# Nodal/Smad2 signaling sustains developmental pausing by repressing Pparg-mediated lipid metabolism

**DOI:** 10.1101/2025.09.08.674904

**Authors:** Giacomo Furlan, S. Bryn Martin, Brandon Cho, Sarah A. McClymont, Elizabeth J. Robertson, Evelyne Collignon, Miguel Ramalho-Santos

## Abstract

Cells and organisms can enter transient dormant states to survive unfavorable conditions during development, physiological adult contexts, and disease. One paradigmatic case of dormancy is diapause, whereby embryos transiently pause development and enter a state of suspended animation. In mammals, diapause occurs pre-implantation at the blastocyst state and involves global growth suppression and metabolic rewiring towards lipid usage as energy source. The molecular regulation of diapause remains poorly understood, including whether it occurs by default or requires active signaling. Here, we identify the Transforming Growth Factor-beta (TGF-β) signaling pathway as an essential driver of transcriptional and metabolic reprogramming in diapause. TGF-β signaling was thought to only be required post-implantation, but we show that the ligand Nodal and its downstream effector Smad2 are essential for the survival of paused embryonic stem cells (ESCs) and blastocysts. Mechanistically, we found that Smad2 represses peroxisome-proliferator activated receptor gamma (Pparg), a transcription factor that is a master regulator of lipid storage, a process incompatible with pausing. Ablation of *Pparg* in *Smad2*-deficient ESCs rescues their survival and prevents excess lipid buildup in paused conditions. Our findings establish Nodal/Smad2 signaling as pivotal for sustaining the transcriptional and metabolic programs of embryonic diapause. This crosstalk between TGF-β signaling and the Pparg pathway may be redeployed in other contexts, such as in cancer dormancy and metabolic disorders.

## Introduction

Embryonic diapause is a reversible developmental arrest occurring at the blastocyst stage in mammals, prior to implantation, in response to environmental cues such as adverse nutritional conditions (Renfree and Fenelon, 2017). In this state, the embryo remains dormant without implanting into the uterine wall. Diapause ends when environmental conditions improve, allowing the embryo to implant and resume active growth in response to specific hormonal cues. This adaptive strategy, observed across many species, enables a better alignment of pregnancy progression with favorable environmental conditions, thereby enhancing offspring survival and overall reproductive success (Garcia-Ojalvo and Bulut-Karslioglu, 2023; Rüegg and Ulbrich, 2023).

In mice, diapaused blastocysts maintain features of pluripotency but display a sharp decrease in proliferation, macromolecular synthesis (including RNA transcription and protein translation), and activity of key growth-promoting pathways, such as Myc and mTOR (Boroviak et al., 2015; Bulut-Karslioglu et al., 2016; Scognamiglio et al., 2016; van der Weijden and Bulut-Karslioglu, 2021). Recent studies indicate that diapause involves large-scale changes in DNA methylation, histone modifications, RNA methylation, and microRNAs (Bulut-Karslioglu et al., 2016; Collignon et al., 2023; Hiratsuka et al., 2023; Iyer et al., 2024; Liu et al., 2020; Stötzel et al., 2024). This state of dormancy also includes deep metabolic rewiring, with a notable decrease in glycolysis and basal respiration concomitant with a shift toward increased fatty acid oxidation as the primary source of energy (Hussein et al., 2020; van der Weijden et al., 2024). Despite these recent insights, it remains unclear whether diapaused blastocysts have specific signaling requirements, and whether and how signaling pathways interact with the chromatin, transcriptional and metabolic reprogramming that underly diapause.

In this study, we first set out to dissect the reprogramming of the chromatin landscape in mouse diapause (see below). The results led us to uncover a pivotal role for the Transforming Growth Factor-beta (TGF-β) pathway during diapause. TGF-β signaling plays crucial roles in the regulation of development, adult stem cells, wound healing and inflammation, and its dysregulation is a major driver of cancer progression (Barcellos-Hoff and Gulley, 2023; Massagué and Sheppard, 2023; Mullen and Wrana, 2017; Wang and Thiery, 2021). Signaling initiates with the binding of TGF-β family ligands (such as TGF-β itself, Nodal or BMPs) to serine/threonine kinase receptors on the cell surface. This binding leads to the downstream phosphorylation and dimerization of Smad proteins. These proteins then form a complex with Smad4, translocate to the nucleus and regulate the expression of target genes (Massagué and Sheppard, 2023; Mullen and Wrana, 2017). TGF-β signaling is essential during early post-implantation development, where it regulates embryonic patterning, gastrulation and mesoderm differentiation (Chiu et al., 2014; Mullen and Wrana, 2017; Robertson, 2014; Senft et al., 2018).

Despite the broad relevance of the TGF-β signaling post-implantation, mouse genetic experiments to date indicate that this pathway is dispensable for pre-implantation development (Datto et al., 1999; Senft et al., 2018; Sirard et al., 1998; Waldrip et al., 1998). In contrast, we uncover here a pivotal role for the TGF-β pathway, and specifically Nodal/Smad2 signaling, in orchestrating the transcriptional and metabolic programs of paused ESCs and blastocysts. Mechanistically, we show that the requirement for Nodal/Smad2 signaling in diapause lies in repressing peroxisome-proliferator activated receptor gamma (Pparg)-mediated lipid storage, thereby maintaining a viable dormant state.

## Results

### Chromatin and transcriptional dynamics highlight TGF-β/Nodal signaling in paused ESCs

To begin uncovering the mechanisms underlying paused pluripotency, we focused on early changes in the chromatin landscape. We performed Assay for Transposase-Accessible Chromatin with high throughput sequencing (ATAC-seq) on mouse ESCs after 24 hours of mTOR inhibition, which we have previously shown captures the diapaused-like state in vitro (Bulut-Karslioglu et al., 2016) (Fig. 1A). In agreement with a previous study (Stötzel et al., 2024), we found that pausing leads to rapid and widespread reprogramming of chromatin accessibility in ESCs (Fig. 1B, S1A-D). Interestingly, most differentially accessible regions after 24h of mTOR inhibition are cases of increased open chromatin (Fig. 1B, S1B). Gene Ontology (GO) analysis of genes in the vicinity of these regions of increased chromatin accessibility during entry into pausing revealed TGF-β signaling as the top-enriched pathway (Fig. 1C-D, S1E). These findings were intriguing, given that TGF-β signalling is largely considered dispensable in pre-implantation pluripotency, in contrast to its known involvement in lineage differentiation at later stages of development (Mullen and Wrana, 2017; Robertson, 2014; Senft et al., 2018). Therefore, we investigated a potential contribution of TGF-β signaling in the regulation of diapause.

**Figure 1.**
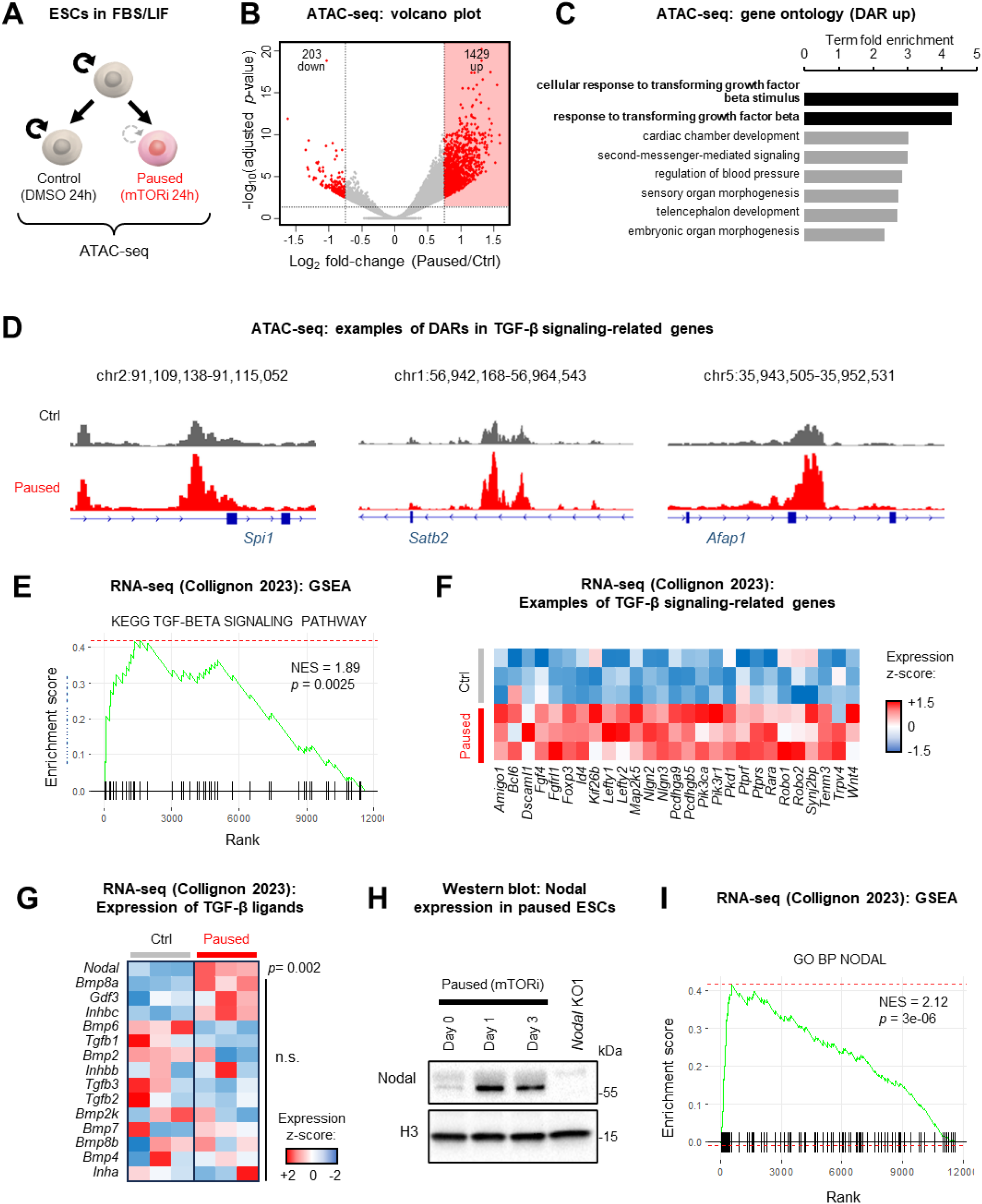
Chromatin and transcriptional dynamics highlight TGF-β/Nodal signaling in paused ESCs. A. Schematic of ESC pausing induced by mTOR inhibition, followed by ATAC-seq (n = 4 biological replicates) B. Volcano plot of ATAC-seq data between paused and control conditions. Significant sites (log_2_ fold-change > 0.75 and False Discovery Rate [FDR] < 0.05) are highlighted in red. C. Gene Ontology analysis of sites with increased accessibility in paused conditions (by ATAC-seq) using Enricher identifies TGF-β signaling as the top regulated pathway D. Examples of ATAC-Seq tracks of TGF-β signaling pathway associated genes enriched in paused conditions. E. GSEA using our previously published RNA-seq data (Collignon et al., 2023) reveals significant enrichment of the ‘KEGG TGF-beta signaling pathway’ in paused ESCs (n = 3 biological replicates). F. Examples of genes related to the TGF-β/Nodal signaling that are upregulated in paused ESCs. G. Expression of TGF-β family ligands in control and paused ESCs using public RNA-seq data (Collignon et al., 2023). H. Western blot analysis of Nodal in control, 1-day, and 3-day paused ESCs. Representative images of n = 2 blots, with histone H3 as loading control and *Nodal* KO1 as negative control. I. GSEA analysis using published RNA-seq data (Collignon et al., 2023) reveals significant enrichment of the ‘GO BP NODAL signaling’ pathway in paused ESCs. P-value by two-sided pre-ranked GSEA with FDR correction (E, I), or two-way paired t-test with FDR correction (G).

We examined RNA-seq data from paused ESCs (Collignon et al., 2023). Gene set enrichment analysis (GSEA) revealed that the TGF-β signaling pathway is significantly enriched at the transcriptional level in paused ESCs (Fig. 1E). Examples of the expression of genes associated with TGF-β pathway activity in control and paused conditions are shown in Fig. 1F. Interestingly, of all potential activating ligands of the TGF-β pathway, only *Nodal* is significantly upregulated in paused ESCs (Fig. 1G). Moreover, Nodal protein is strongly induced upon transition to the paused state (Fig. 1H). In line with these results, a GO signature of the Nodal pathway is enriched in paused ESCs (Fig. 1I). Taken together, these data suggest that the Nodal branch of the TGF-β pathway is active and may play a pivotal role in chromatin and transcriptional dynamics in ESCs upon entry into the paused state.

### The Nodal/Smad2 signaling is essential for survival of ESCs specifically in the paused state

We next tested a potential role of Nodal in paused pluripotency, using two independent clones of *Nodal* KO ESCs (Varlet et al., 1997). Proliferation assays revealed a striking vulnerability of *Nodal* KO ESCs in paused conditions, marked by a significant reduction in cell numbers (Fig. 2A). Similar results were obtained using SB431542, a selective chemical inhibitor of the Nodal branch of the TGF-β pathway at the receptor level (Fig. S2A) (Du et al., 2014). In paused conditions, inhibition of Nodal signaling induced a sharp reduction in cell proliferation, whereas this treatment had no impact in control unpaused conditions (Fig. S2A). Large numbers of dying cells were observed in *Nodal* KO ESCs upon entry into the paused state, a finding quantified using flow cytometry measurements of apoptosis (Fig.2B), whereas wild-type (WT) paused ESCs were largely unaffected (Fig. 2B). In contrast, no significant difference in viability was observed under control culture conditions, indicating that Nodal is specifically required to support cell survival during paused pluripotency (Fig. S2B).

**Figure 2.**
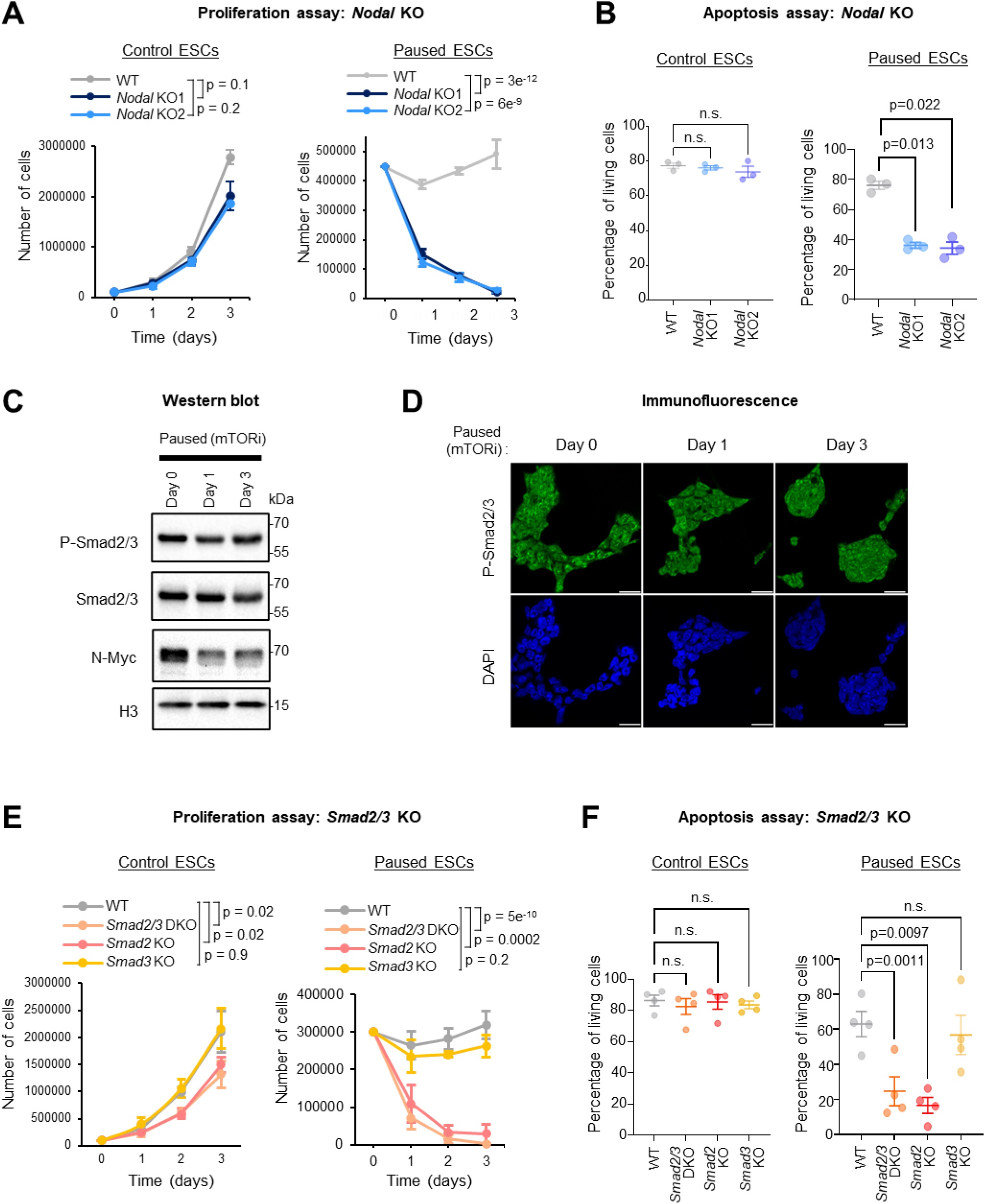
The Nodal/Smad2 signaling is essential for survival of ESCs specifically in the paused state. A. Proliferation assay of WT and two clones of *Nodal* KO ESCs in control (left) or paused (right) conditions. Paused *Nodal* KO ESCs show a strong decrease in cell numbers. Data are presented as mean ± SD (n =3 biological replicates). B. Apoptosis assay by flow cytometry using PI/Annexin staining in control (left) and paused (right) ESCs, showing reduced viability in paused *Nodal* KO1 and KO2 ESCs (n = 3 biological replicates). C. Western blot analysis of phospho-Smad2/3, total Smad2/3, and N-Myc in control, 1-day, and 3-day paused ESCs. Representative images of n = 2 blots, with histone H3 as loading control. D. Immunofluorescence staining of phospho-Smad2/3 in control or 1-day and 3-day paused ESCs. Representative images of n = 2 biological replicates, with DAPI (FxCycle Violet Stain) for nuclear staining. E. Proliferation assay of WT, *Smad2/3* KO, *Smad2* KO, and *Smad3* KO ESCs in control (left) or paused conditions (right). Paused *Smad2/3* DKO and *Smad2* KO ESCs show a strong decrease in cell numbers. Data are presented as mean ± SD (n =3 biological replicates). F. Apoptosis assay by flow cytometry using PI/Annexin staining in control (left) or paused conditions (right), showing reduced viability in paused *Smad2/3* DKO and *Smad2* KO ESCs (n = 4 biological replicates). P-values by linear regression test with interaction (A, E), or one-way analysis of variance (ANOVA) with Dunnett’s multiple comparison test (B, F).

Activation of the Nodal branch of the TGF-β pathway leads to phosphorylation of the downstream effectors Smad2/3 (Massagué and Sheppard, 2023; Mullen and Wrana, 2017). Western blotting and immunofluorescence confirmed that this branch, which is known to be active in control ESCs (Senft et al., 2018), remains active in the paused state (Fig. 2C-D). This persistence of TGF-β/Nodal signaling in paused ESCs stands in contrast to the downregulation of most molecular pathways in this state (Bulut-Karslioglu et al., 2016; Scognamiglio et al., 2016), illustrated here by the decreased level of N-Myc (Fig. 2C). We therefore explored a potential role for Smad2 and Smad3 in paused ESCs. We began by using *Smad2/3* double knockout (DKO) ESCs (Senft et al., 2018). The proliferation phenotype of *Smad2/3* DKO ESCs closely resembles that of *Nodal* KO cells, with a marked reduction in cell number specifically in paused conditions (Fig. 2E). Analyses of individual *Smad2* or *Smad3* KO ESCs revealed that this phenotype is primarily driven by the loss of *Smad2* (Fig. 2E, S2C). Apoptosis analyses confirmed that the drastic drop in cell numbers in *Smad2/3* DKO and *Smad2* KO paused ESCs is due to a decrease in cell survival (Fig. 2F, S2D). Here again, no differences in cell viability were observed in the Smad mutant ESCs in control culture conditions (Fig. 2F).

Taken together, these results reveal that the Nodal branch of the TGF-β pathway, and specifically the Nodal ligand and Smad2 effector, is essential to sustain ESCs in the paused pluripotent state. While negligible under standard culture conditions, Nodal/Smad2 signaling disruption becomes synthetically lethal with the induction of the paused state in ESCs.

### The Nodal/Smad2 axis is essential for embryonic diapause

We next sought to explore the dynamics of Nodal signaling during diapause in vivo. We analyzed a single-cell RNA sequencing (scRNA-seq) dataset comprising control blastocysts at E4.5 and diapaused blastocysts at equivalent day of gestation (EDG) 7.5 and 9.5 (Chen et al., 2024). In agreement with previous findings (Granier et al., 2011; Mesnard et al., 2006), *Nodal* is detected primarily in epiblast cells, with much lower level expression also detected in some primitive endoderm cells (Fig. 3A, S3A). By focusing on the epiblast compartment, we found that the expression of *Nodal* and its co-receptor Cripto is significantly increased in diapaused embryos (Fig. 3A-B, S3B-C). Moreover, the GO signature of the Nodal pathway is also significantly enriched in the diapaused epiblast, compared to E4.5 controls (Fig. 3C). The activation of Nodal expression and signaling in diapaused blastocysts mirrors the results in paused ESCs (Fig. 1G-I) and stands in contrast with the known repression of many other pathways during embryonic pausing (Bulut-Karslioglu et al., 2016) (e.g., Myc, mTORC1, oxidative phosphorylation, see Fig. S3D). To further validate these observations, we compiled bulk RNA-seq data from various studies, confirming the upregulation of *Nodal* in diapaused embryos (Fig. S3E).

**Figure 3.**
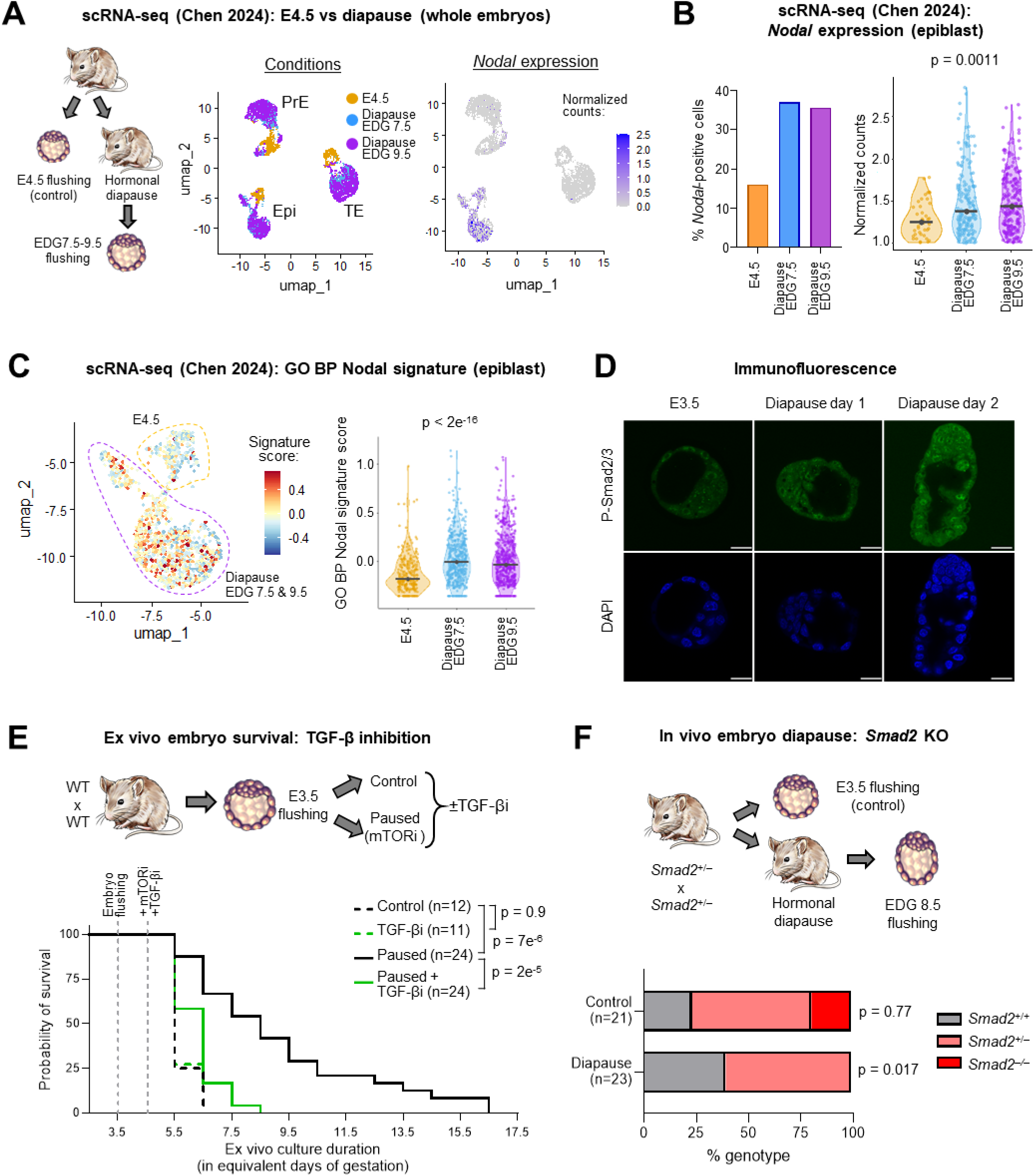
The Nodal/Smad2 axis is essential for embryonic diapause. A. Public single-cell RNA-seq data (Chen et al., 2024) reveal that *Nodal* expression is mostly confined to the epiblast cells, in both diapaused (equivalent days of gestation [EDG] 7.5 and 9.5) and control embryos (E4.5). B. The percentage of *Nodal*-positive cells (left) and the level of *Nodal* expression among them (right) are increased in diapaused epiblast cells. C. Zoomed in view of UMAP in Fig. 3A focusing on Nodal expression in epiblast cells reveals increased ‘GO BP Nodal’ signature score in the epiblast cells of diapaused embryos. D. Immunofluorescence staining of phospho-Smad2/3 in diapaused and control mouse blastocysts. Representative images of n = 2 biological replicates, with DAPI (FxCycle Violet Stain) for nuclear staining. E. Kaplan-Meyer curve analysis depicting ex vivo survival of embryos treated or not with 10µM TGF-β signaling inhibitor SB431542, in control and pausing conditions. TGF-β inhibition leads to premature death of mouse blastocysts cultured ex vivo in paused conditions. Number of embryos (n) as indicated. F. Genotype distribution of recovered (live) embryos resulting from *Smad2*^+/−^ crossing, with embryos collected either at E3.5 (control) or at EDG 8.5 following hormonal diapause, showing that *Smad2*^−/−^ embryos are impaired at undergoing hormonal diapause. Number of embryos (*n*) as indicated. P-values by one-way ANOVA (B, C), log-rank test (E), or *χ*^2^ test against expected Mendelian ratios (F).

To explore a role for Nodal signaling in embryonic diapause, we first performed immunofluorescence on embryos upon induction of hormonal diapause. We found that phospho-Smad2/3 levels are maintained in diapaused blastocysts, indicating that the pathway remains active (Fig. 3D), consistent with the case in paused ESCs (Fig. 2C-D). We then tested the effects of the inhibition of Nodal/Smad2-mediated signaling using two embryo models. In the first model, we carried out mTOR inhibition in ex vivo cultures of blastocysts, which we have shown induces a diapause-like state (Bulut-Karslioglu et al., 2016; Collignon et al., 2023). Using this model, paused blastocysts survive for up to two weeks ex vivo (Fig. 3E). However, treatment with the SB431542 inhibitor significantly reduces blastocyst survival, leading to complete embryo loss by EDG 8.5 (Fig. 3E). In the second model, we intercrossed *Smad2* heterozygous KO mice (*Smad2*^+/−)^ and hormonally induced diapause in pregnant females (MacLean Hunter and Evans, 1999; Weitlauf and Greenwald, 1968). *Smad2*^−/−^ blastocysts are recovered at the expected Mendelian frequency when embryos are collected at E3.5 without induction of diapause, in agreement with previous reports that Smad2 is dispensable for pre-implantation development (Waldrip et al., 1998) (Fig. 3F, S3F). In contrast, we found that hormonal induction of diapause leads to a complete loss of *Smad2*^−/−^ embryos by EDG

8.5 (Fig. 3F). Thus, similarly to the case in paused ESCs, canonical TGF-β signaling via the Nodal/Smad2 axis is strictly and specifically required for the survival of diapaused embryos.

### Smad2 orchestrates gene expression and lipid metabolism in paused ESCs

To dissect the molecular mechanisms of Smad2-mediated signaling in the paused pluripotent state, we generated additional clones of *Smad2* KO ESCs using a CRISPR-Cas9 approach (Fig. 4A). We validated that these *Smad2* KO ESCs display complete loss of Smad2 protein (Fig. S4A) and proliferation defects in the paused state (Fig. S4B). We used these *Smad2* KO ESCs to perform RNA-Seq in both control and paused conditions (Fig. 4B-C). Samples were collected at 12 hours of induction of pausing by mTOR inhibition, to capture early transcriptional changes and avoid secondary effects from cell death. Principal component analysis (PCA) and hierarchical clustering revealed a clear separation of samples by both culture condition and by genotype (Fig. 4B-C), suggesting that Smad2 loss reshapes the transcriptome in both control and paused ESCs, without affecting the expression of key pluripotency regulators (Fig. 4B-C, S4D). The two clones of *Smad2* KO ESCs displayed transcriptomes highly similar to each other and clearly distinct from wild-type (WT) cells (Fig. 4B-C, S4C); they were therefore treated as replicates in subsequent analyses.

**Figure 4.**
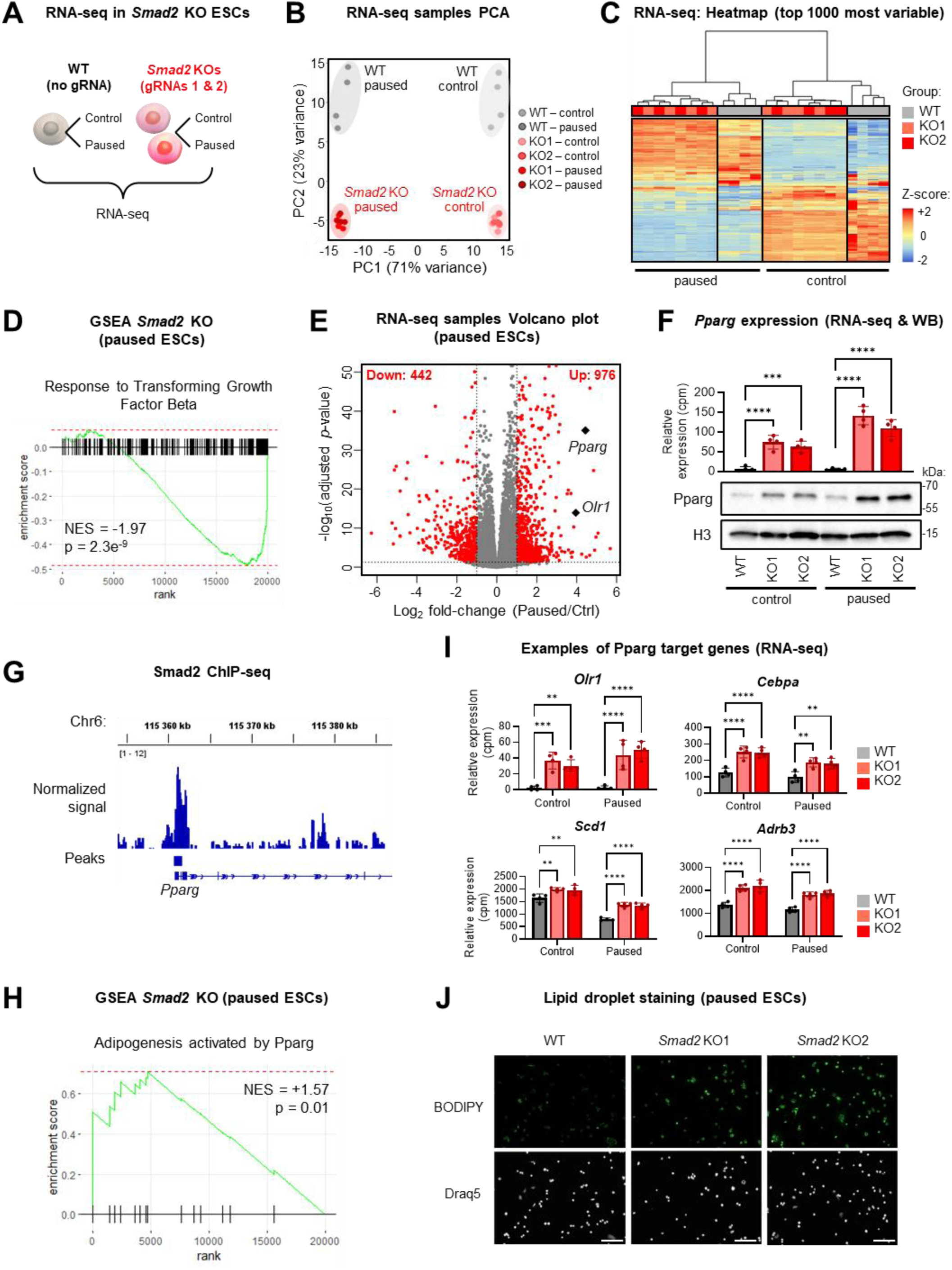
Smad2 orchestrates gene expression and lipid metabolism in paused ESCs. A. Schematic of RNA-seq experiment. WT and in-house *Smad2* knockout ESCs (generated with *Smad2*-targeting gRNAs 1 & 2) were cultured in control and paused conditions, followed by RNA-seq. B. PCA plot of RNA-seq data segregates ESCs according to pausing state condition (along PC1) and genotype (along PC2). *Smad2* KO1 and KO2 display highly consistent transcriptional profiles. N = 4 biological replicates per group. C. Heatmap of the top 1000 most variable genes, identified by RNA-seq, with unsupervised hierarchical clustering. D. GSEA on RNA-seq data reveals reduced TGF-β signaling in *Smad2* KO paused ESCs (aggregated from KO1 and KO2). E. Volcano plot of RNA-seq data between *Smad2* KO and WT paused conditions. Significant genes (log_2_ fold-change > 1 and FDR < 0.05) are highlighted in red. *Pparg* and its downstream target *Olr1* are among the top upregulated genes. E. F. *Pparg* is upregulated in *Smad2* KO ESCs, in both control and paused conditions, as assessed by RNA-seq (top) and western blot (bottom). Representative images of n = 3 blots, with histone H3 as loading control. G. Published ChIP-seq data reveal enriched binding of Smad2 at the promoter of the *Pparg* gene in ESCs. Data were aggregated from several public ChIP-seq (n = 7, see methods) (GSE240787, GSE186222, GSM5983538). H-I. Increased activity of Pparg in paused *Smad2* KO ESCs, as assessed by enrichment of the ‘Adipogenesis activated by Pparg’ gene signature (H, by GSEA on RNA-seq data), with examples of upregulated Pparg target genes shown (I). J. BODIPY live staining by immunofluorescence shows increased lipid accumulation in paused *Smad2* KO ESCs. Nuclei were stained with Draq5 as control. Representative images of n = 3 independent replicates. P-values by two-sided pre-ranked GSEA (D, H), or two-way ANOVA with Dunnett’s multiple comparison test (F top, I).

As expected, *Smad2* KO paused ESCs display a marked reduction in the transcriptomic signature of the TGF-β pathway (Fig. 4D, S4E). Among the upregulated signatures in *Smad2* KO paused ESCs are several pathways associated with lipid metabolism, notably lipid storage (Fig. S4E). This result is of interest because lipid breakdown by fatty acid oxidation has recently been shown to be essential for sustaining mammalian embryos during diapause (Arena et al., 2021; van der Weijden et al., 2024). In agreement with these findings, *Pparg* and its downstream target *Oxidized low-density lipoprotein receptor 1 (Olr1)* are among the top upregulated genes in *Smad2* KO paused ESCs (Fig. 4E-F). Pparg is a nuclear receptor that plays a central role in lipid metabolism and adipogenesis (Hernandez-Quiles et al., 2021; Sun et al., 2024). It has been shown to mediate lipid dynamics during trophectoderm and placenta development (Fournier et al., 2008; McGraw et al., 2024; Ribeiro et al., 2016), but has not been implicated in the regulation of pluripotency or diapause.

We decided to explore further a potential role for Pparg in disrupting the paused state of *Smad2* KO ESCs. Pparg is expressed at negligible levels in wildtype ESCs, both in paused and control conditions, but it is highly upregulated at the mRNA and protein level in *Smad2* KO ESCs, particularly in the paused state (Fig. 4F). Interestingly, our re-analysis of published data for Smad2 ChIP-seq in ESCs (Baas et al., 2016; Sun et al., 2023; Zhao et al., 2024) revealed that Smad2 binds to the promoter of the *Pparg* gene (Fig. 4G). Together with the strong induction of Pparg in Smad2 KO ESCs, these results suggest that Smad2 may directly repress *Pparg* at the transcriptional level. In line with these observations, a signature of adipogenesis genes driven by Pparg is strongly induced in *Smad2* KO paused ESCs (Fig. 4H). Examples of direct Pparg target genes that mediate lipid anabolism and are upregulated in *Smad2* KO ESCs are shown in Fig. 4I. These data indicate that a Pparg-mediated program of lipid biosynthesis and storage is abnormally activated upon loss of *Smad2* in ESCs. In support of this notion, we observed an aberrant accumulation of lipid droplets in *Smad2* KO paused ESCs (Fig. 4J). Collectively, these findings indicate that Smad2 is required to suppress a Pparg-driven program of lipid storage in paused ESCs.

### Smad2 safeguards against a Pparg-driven chromatin, transcriptional and metabolic program incompatible with the paused state

Given the extensive transcriptional rewiring observed in *Smad2* KO ESCs, we sought to gain further mechanistic insights by investigating the landscape of chromatin accessibility. We performed ATAC-seq on *Smad2* KO and WT ESCs in control and pausing conditions. PCA and hierarchical clustering revealed that loss of *Smad2* induces a significant reprogramming of chromatin accessibility in ESCs (Fig. 5A-C), consistent with the RNA-seq data (Fig. 4B-C). Moreover, there is an overall positive correlation between the ATAC-seq and RNA-seq datasets, wherein genes showing increased accessibility tend to be upregulated at the transcriptional level, and vice-versa (Fig. S5A-B).

**Figure 5.**
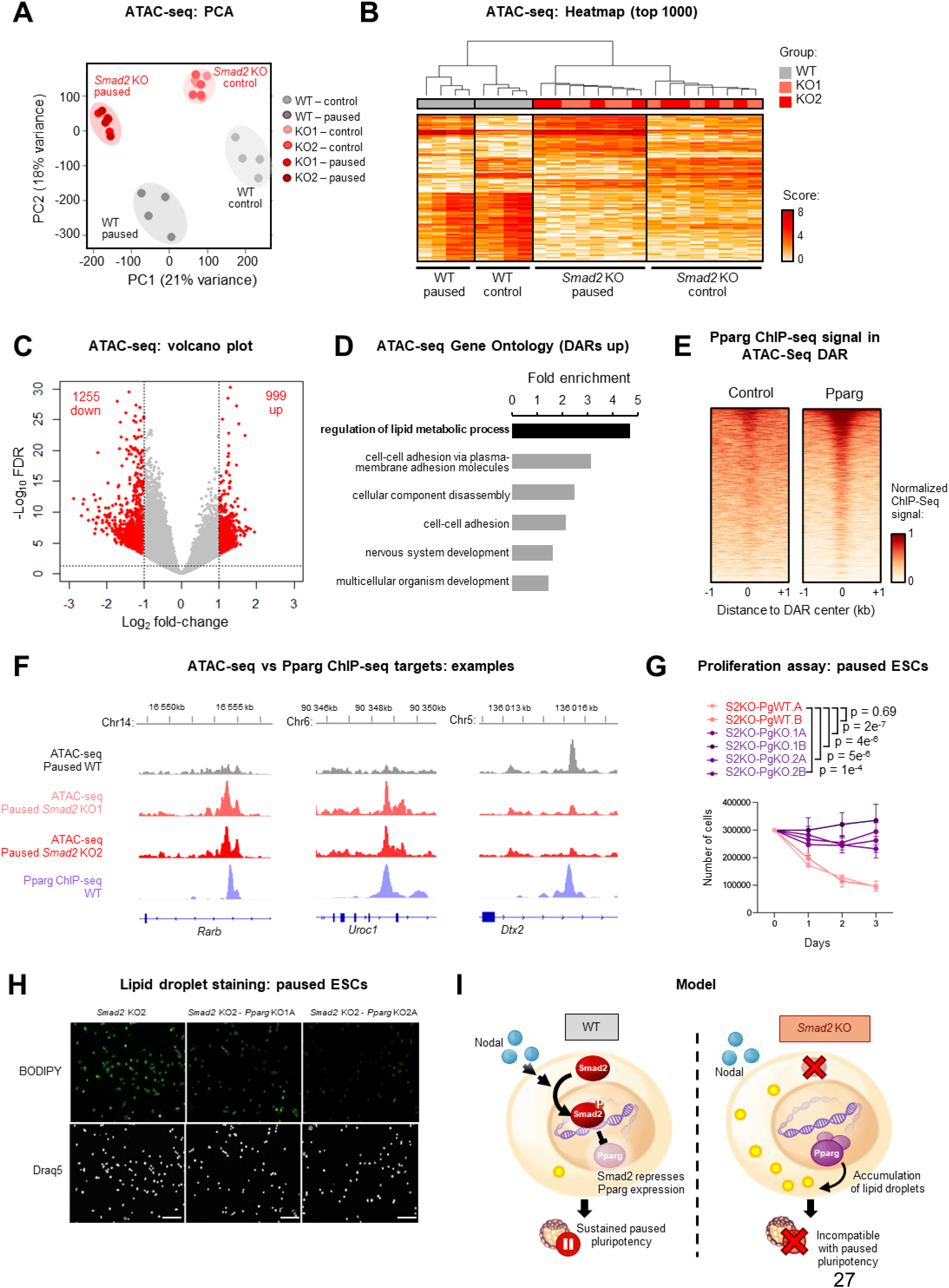
Smad2 safeguards against a Pparg-driven chromatin, transcriptional and metabolic program incompatible with the paused state. A. PCA plot of ATAC-seq data segregates ESCs according to paused state condition and genotype. *Smad2* KO1 and KO2 display highly similar profiles. N = 4 biological replicates per group. B. Heatmap of the top 1000 most variable genes, identified by ATAC-seq, with unsupervised hierarchical clustering. C. Volcano plot of RNA-seq data between *Smad2* KO and WT paused conditions. Significant sites (log_2_ fold-change > 1 and FDR < 0.05) are highlighted in red. D. Top pathways enriched among sites with increased accessibility (by ATAC-seq) in paused *Smad2* KO using Panther Gene Ontology. E-F. Heatmap of Pparg ChIP-seq (via HA-tagged Pparg) and control (HA-only) signals, showing enrichment of Pparg among sites with increased genome accessibility in paused *Smad2* KO ESCs (E), with examples of ATAC-seq and ChIP-seq tracks shown (F). Pparg ChIP-seq data were aggregated from n = 2 independent replicates, using public data (GSE199426) (Madsen et al., 2022). G. Proliferation assay showing reduced cell numbers in paused *Smad2* KO (S2KO-PgWT) ESCs, consistent with the previously observed pausing defect (see Fig. 2A). This phenotype is rescued in *Smad2–Pparg* DKO (S2KO-PgKO) cells (n =3 biological replicates). H. *Smad2*-*Pparg* DKO rescues increased lipid accumulation observed in *Smad2* KO ESCs, as measured by BODIPY staining. Nuclei were stained with Draq5 as a control. Representative images of n = 3 independent replicates. I. Model recapitulating main findings of the study. P-values by linear regression test with interaction (G).

Interestingly, GO analysis of regions with gain in chromatin accessibility in paused *Smad2* KO ESCs identified lipid metabolism as the top enriched pathway (Fig. 5D). ChIP Enrichment Analysis (ChEA) revealed that Pparg is among the top enriched transcription factors that bind these regions (Fig. S5C). To further explore this result, and given that Pparg expression is almost absent in WT ESC (Fig. 4F), we took advantage of a ChIP-seq dataset for Pparg in mouse embryonic fibroblasts (Madsen et al., 2022). This analysis revealed a striking overlap between Pparg binding sites and Smad2-dependent regions of chromatin accessibility (Fig. 5E-F, S5D). Here again, we observed a positive correlation between changes in chromatin accessibility and gene expression at Pparg targets in *Smad2* KO ESCs (Fig. S5E). This convergence suggests that a substantial portion of the chromatin and transcriptional rewiring induced by loss of *Smad2* in paused ESCs is associated with increased Pparg activity.

The results above led us to hypothesize that the upregulation of Pparg may in part mediate the lethality of *Smad2* KO ESCs in the paused state, possibly by disrupting lipid metabolism. To test this, we generated *Smad2*-*Pparg* DKO ESCs by targeting *Pparg* in *Smad2* KO ESCs using CRISPR-Cas9 technology (Fig. S5F). The absence of Pparg protein in DKO ESCs was validated by western blotting (Fig. S5G). Remarkably, genetic ablation of *Pparg* in four independent clones effectively rescued the proliferation defects of *Smad2* KO ESCs in the paused state (Fig. 5G, 2E, S4B). These results indicate that the sharp cell loss observed in paused *Smad2* KO ESCs (Fig. 2E) is driven by the aberrant upregulation of Pparg. In agreement with these findings, the abnormal accumulation of lipid droplets seen in *Smad2* KO paused ESCs is suppressed in *Smad2*-*Pparg* DKO cells (Fig. 5H). Taken together, these data establish that Smad2 is critical for maintaining cell viability during paused pluripotency by repressing a Pparg-driven transcriptional program that drives aberrant lipid storage (Fig. 5I).

## Discussion

In summary, our findings reveal an essential role for TGF-β signaling in the survival of ESCs and blastocysts during embryonic diapause. We found that Nodal is the upstream ligand and Smad2 is the downstream signaling effector that enforces the transcriptional program of developmental pausing. In particular, we found that Smad2 is key to repress Pparg, a master regulator of lipid metabolism. Loss of Pparg repression in Smad2 KO ESCs leads to lipid droplet accumulation and cell death, underscoring the indispensable role of Nodal/Smad2 signaling in embryonic diapause.

These findings highlight a surprising requirement for TGF-β signaling in embryonic diapause. While TGF-β signaling is well-established for its roles in developmental patterning and cell fate, it has largely been considered dispensable for pre-implantation development (Massagué and Sheppard, 2023; Mullen and Wrana, 2017). Notably, genetic ablations of *Smad2* or *Nodal* result in viable ESCs and embryos that can progress to post-implantation developmental stages (Lowe et al., 2001; Senft et al., 2018; Tremblay et al., 2000; Varlet et al., 1997; Waldrip et al., 1998). Nodal is present in the uterus of pregnant mice, synthetized by the glandular epithelium of the endometrium, and its absence leads to placentation defects and fetal loss (Park et al., 2012; Park and Dufort, 2011). It may prove interesting to dissect the role of maternal versus embryonic Nodal in diapause. Mechanistically, the critical role of Nodal/Smad2 in paused pluripotency is linked to the regulation of lipid metabolism, which recently emerged as a cornerstone of diapause. Indeed, lipid catabolism serves as the primary energy source to sustain the paused state (Arena et al., 2021; van der Weijden et al., 2024). Our work reveals that the Nodal/Smad2 pathway sustains developmental pausing by repressing Pparg-driven lipid storage. Interestingly, Nodal/Smad2 signaling is active and represses *Pparg* in control (non-paused) ESCs but only becomes essential for survival upon induction of pausing. This may be because it is only in the paused state that lipids become the primary energy source, as recently reported (van der Weijden et al., 2024). Taken together, our findings provide new insights into the interplay between signaling and metabolic reprogramming during diapause and raise several avenues for future inquiry.

It will be of interest to dissect further the repression of *Pparg* expression by Smad2 during embryonic diapause. For example, *Pparg* expression can be regulated by promoter DNA methylation (Fujiki et al., 2009; Sabatino et al., 2012), and Smad2/3 were recently shown to interact with the DNA methyltransferase Dnmt3b during naïve-to-primed pluripotency and differentiation (Zhao et al., 2024). However, DNA methylation is thought to be dispensable for developmental pausing (Stötzel et al., 2024). Alternatively, Smad2 may repress Pparg by competing for binding of activating transcription factors and/or recruiting co-repressors that enforce silencing histone marks (Lakshmi et al., 2017; Massagué and Sheppard, 2023; Mullen and Wrana, 2017). An additional, non-mutually exclusive possibility is repression of *Pparg* via N6-methyladenosine (m6A) RNA methylation, a layer of gene regulation that we have shown is required for diapause (Collignon et al., 2023). Of note, TGF-β signaling interacts with the RNA methylation machinery in human ESCs (Bertero et al., 2018). Thus, it is possible that multiple functions of Smad2 and its interacting partners contribute to enforce repression of *Pparg* during diapause.

Our findings on an essential function for TGF-β signaling in diapause raises intriguing parallels to the role of this pathway in cancer. Although TGF-β signaling has a context-dependent effect in tumorigenesis, it is generally associated with suppression of cell proliferation and induction of cytostasis (Deng et al., 2024; Gomis et al., 2006; Massagué and Sheppard, 2023), which closely mirrors its function in the dormant state of embryonic diapause reported here. This parallel to cancer is particularly evident in the context of drug-tolerant persister cells (DTPs), a pool of cancer cells that temporarily switch to a dormant state of low proliferation and reduced metabolic activity to evade anti-cancer treatments (Collignon, 2024; Russo et al., 2024). We and others recently reported that the DTP state shares substantial transcriptional and functional similarities with diapause, suggesting that cancer cells co-opt pathways of developmental pausing to evade chemotherapy (Dhimolea et al., 2021; Rehman et al., 2021). In further support of this notion, TGF-β pathway activation promotes the DTP state in various cancers, while its inhibition was shown to increase chemotherapy sensitivity in persister cells (Brown et al., 2017; Marsolier et al., 2022; Moghal et al., 2023; Nonninger et al., 2025; Park et al., 2015). Thus, our findings that Nodal/Smad2 signaling is critical for paused pluripotency underscore a broader role for the TGF-β pathway in promoting dormancy to survive diverse stress conditions. Moreover, our results suggest that manipulating Pparg activity and lipid metabolism may be an additional strategy to specifically target DTP cells and prevent cancer recurrence.

Our results prompt exploration of the connection between the TGF-β pathway and Pparg in the context of the regulation of lipid metabolism more generally. TGF-β signaling regulates adipocyte differentiation and is associated with metabolic disorders such as obesity and diabetes (Choy and Derynck, 2003; Kumari et al., 2021; Tan et al., 2012; Yadav et al., 2011). Of note, Smad3 has been shown to repress *PGC1a* in adipocytes, and PGC1a (Yadav et al., 2011) is a transcriptional regulator and interacting partner of Pparg (Finck, 2006; Gross et al., 2017). It will be of interest to determine whether TGF-β signaling, mediated by Smad2 or Smad3 (Aragón et al., 2019), directly modulates lipid metabolism by repressing Pparg during fat development or homeostasis. Moreover, our data suggest that there may be distinct requirements for TGF-β signaling in lipid metabolism in situations of quiescence, such as possibly in brown fat dormancy (Kutyavin and Chawla, 2019).

In conclusion, our work reveals an essential role for the Nodal/Smad2 axis in embryonic diapause and establishes a critical connection between environmental stress, signaling, metabolic reprogramming and developmental timing. These findings provide new insights into the fundamental biology of dormant states, with broad potential implications for reproductive health, regenerative medicine, cancer recurrence and metabolic disorders.

## Methods

All mouse procedures were carried out in accordance with the *Animals for Research Act* of Ontario and the guidelines set forth by the Canadian Council on Animal Care. Experimental protocols were reviewed and approved by the Animal Care Committee at The Centre for Phenogenomics, Toronto (TCP, protocol 26-0331H).

### Mouse models

Mice were housed in a pathogen-free environment at the TCP, in individually ventilated cages (Tecniplast) under controlled conditions (18–23 °C, 40–60% humidity) and a 12-hour light/dark cycle, with food and water provided *ad libitum*.

The B6;129-Smad2^tm1Rob^ mouse line was kindly donated by the E. Robertson lab (Waldrip et al., 1998), received as sperm and rederived onto a C57BL/6J background at the in-house TCP facility. Heterozygote mice were maintained on a C57BL/6J background and genotyped as previously described (Waldrip et al., 1998).

### Hormonal diapause experiments

For hormonal diapause experiments, heterozygous Smad2^tm1Rob^ C57BL/6J males were crossed with CD1 wild-type females to generate C57BL/6J × CD1 hybrid offspring, which are more suitable for diapause studies than pure C57BL/6J mice. At 8–12 weeks of age, heterozygous Smad2^tm1Rob^ C57BL/6J × CD1 hybrid mice (hereafter referred to as Smad2^+/-^) were intercrossed, and females were checked the following morning for the presence of a vaginal plug (designated as embryonic day 0.5, E0.5).

At E2.5, pregnant females were injected subcutaneously with 100μL of a 30mg/mL medroxyprogesterone 17-acetate solution (Sigma, #M1629-1G), and intraperitoneally with 100μL of a 100μg/mL tamoxifen solution (Sigma, #T5648). Both hormones were dissolved in peanut oil. For six days, from E2.5 to EDG8.5 (equivalent day of gestation 8.5), females received daily subcutaneous injections of medroxyprogesterone 17-acetate. A single additional tamoxifen injection was administered at EDG5.5. At EDG8.5, females were euthanized and embryos collected for subsequent analyses and genotyping. Blastocysts with a completely collapsed blastocoel were considered non-viable and excluded from genotyping.

Genomic DNA was extracted from individual blastocysts using the PicoPure™ DNA Extraction Kit (Life Technologies, #KIT0103) following the manufacturer’s instructions. PCR genotyping was performed using the Phire Green Hot Start II PCR Master Mix (Life Technologies, #F126L) with the following thermal cycling conditions: 95 °C for 3 min; 35 cycles of 95 °C for 30 sec, 65 °C for 1 min, 72 °C for 1 min; and a final extension at 72 °C for 5 min. PCR primer sequences are listed in Table 1.

### Ex vivo pausing experiments

Collection and ex vivo culture of blastocysts were carried out as previously described (Bulut-Karslioglu et al., 2016). Briefly, 8-12 week old CD1 wild-type males and females were mated, and plugged females were singly housed and sacrificed at E3.5. Blastocysts were collected by uterine flushing using EmbryoMax M2 Medium (Sigma, #MR-015-D) and cultured at 37 °C in a 5% O₂, 5% CO₂ incubator using EmbryoMax Advanced KSOM Embryo Medium (Sigma, #MR-101-D). One day after flushing (E4.5), 200nM mTOR inhibitor RapaLink-1 (MedchemExpress, #HY-111373-1MG) was added to induce pausing ex vivo. To inhibit TGF-β signaling, 10μM TGF-βRI kinase inhibitor SB431542 (Sigma, #616464-5MG) was also added to the culture medium. To generate survival curves, viable embryos were examined daily.

### Mouse ES cell culture and treatments

Mouse embryonic stem cells (mESCs) were cultured on gelatin-coated plates (Sigma, #G1393) in standard serum/LIF medium consisting of DMEM GlutaMAX supplemented with sodium pyruvate (Life Technologies, #10569044), 15% fetal bovine serum (BioTechne, #S12450), 0.1 mM non-essential amino acids (Life Technologies, #11140050), 50 U mL^−1^ penicillin–streptomycin (Life Technologies, #15140122), 0.1 mM 2-mercaptoethanol (Fisher Scientific, # 21985023), and 1,000 U mL^−1^ ESGRO LIF (produced in-house).

To induce in vitro pausing, 200nM of the mTOR inhibitor INK128 (AbMole, #M2151) was added to the culture medium, as previously described (Bulut-Karslioglu et al., 2016). Unless otherwise stated, mESCs were maintained in serum/LIF and paused for 12 or 24 hours for sequencing experiments), 48 hours for apoptosis assays, and 72 hours for proliferation curve analyses, as detailed in the corresponding figure legends.

To inhibit TGF-β signaling, 10μM SB431542 (Sigma, #616464-5MG) was also added to the culture medium.

### Cell models and cloning

*Smad2* knockout (KO), *Smad3* KO, *Smad2/3* double knockout (DKO) and *Nodal* KO mESC lines were provided by E. Robertson lab (Sir William Dunn School of Pathology) and derived as previously described (Senft et al., 2018; Tremblay et al., 2000; Varlet et al., 1997).

Independent Smad2 KO1 and KO2 mESC lines were generated in-house using CRISPR-Cas9 deletion techniques. Guide RNAs targeting *Smad2* were cloned into the pSpCas9(BB)-2A-GFP (PX458-GFP) plasmid, a gift from F. Zhang (Addgene plasmid #48138) (Ran et al., 2013). Two distinct guide sets were used to generate the KO1 and KO2 lines. E14 mESCs were transfected with the constructed plasmids using Lipofectamine 3000 (Life Technologies, #L3000015), sorted by FACS for GFP-positive cells, and seeded as single cells to derive clonal lines. WT clones were transfected with empty PX458-GFP, while KO1 and KO2 clones were transfected with the corresponding *Smad2*-targeting plasmids. Following expansion, clonal lines were validated for *Smad2* KO by western blot.

*Smad2*-*Pparg* double knockout (DKO) lines were derived using the same CRISPR–Cas9 strategy. Guide RNAs targeting *Pparg* were cloned into PX458-BFP and transfected into *Smad2* KO2 clonal line. BFP-positive cells were FACS-sorted, seeded as single cells, expanded, and validated for western blot.

### Cell phenotypical assays

For proliferation curve analyses, 100,000 cells were plated per well for control conditions and 300,000 cells per well for paused conditions. Six hours after plating, cells in the paused condition were treated with 200nM INK128 to induce pausing. Cells were harvested and counted daily for three consecutive days using the Bio-Rad TC20™ Automated Cell Counter.

For apoptosis assays, cells were plated and treated identically to the proliferation experiments. 48 hours after treatment, cells were collected and stained in 250μl of Annexin V Binding Buffer (BioLegend, #422201) containing 0.3μL of Annexin V-Alexa Fluor 647 conjugate (Invitrogen, #A23204) and 0.25μg of propidium iodide (PI, Sigma, #P4170-10MG) for 15 minutes. Stained cells were analyzed by flow cytometry using a Gallios Flow Cytometer.

### RNA-seq library preparation and data analysis

For each sample, total RNA was extracted from 50,000 cells using the RNeasy Mini Kit (Qiagen, #74106). For library preparation, 100ng of RNA per sample was processed using the NEBNext Ultra II Directional RNA Library Prep Kit for Illumina (NEB, #E7760L), in combination with the NEBNext Poly(A) mRNA Magnetic Isolation Module (NEB, #E7490S), according to the manufacturer’s instructions. Sequencing was performed at The Centre for Applied Genomics (TCAG) at SickKids (Toronto) on a NovaSeq X platform with a 10B flow cell, generating 150 bp single-end reads.

For data analysis, raw libraries underwent quality control and adapter trimming using FastQC (v0.12.1) and Trim Galore (v0.6.6). Reads were aligned to the *Mus musculus* (mm10) reference genome using HISAT2 (v2.2.1). Raw counts per gene were obtained using the featureCounts function from Rsubread (v2.20.0) with parameters -t exon -g gene_id. Raw counts were imported into R (v4.4.1) and normalized using DESeq2 (v1.46.0). Differential gene expression was considered significant with an adjusted p-value (Padj) < 0.05 and an absolute log₂ fold change > 1. Counts per million (CPM) were calculated using the cpm function in edgeR (v3.36.0). Data visualization was performed using ggplot2 (v3.4.1) or GraphPad Prism (v.9.3.1). Principal component analysis (PCA) and heatmaps were generated from normalized count data using the DESeq2 and pheatmap (v1.0.12) packages. Gene ontology (GO) enrichment analysis was performed using the PANTHER database (v19.0), with background genes defined as those expressed in at least one of the conditions (WT or *Smad2* KO, in control or paused states). GSEA was carried out with the fGSEA package (v.1.26.0), with genes pre-ranked by fold-change from the differential analysis (paused/control or paused *Smad2* KO/paused WT). Gene set collections were downloaded from the Molecular Signatures Database (v.7.5.1; http://www.gsea-msigdb.org/gsea/msigdb/index.jsp).

### ATAC-Seq library preparation and data analysis

ATAC-seq libraries were prepared following the Omni-ATAC protocol (Corces et al., 2017). Briefly, 50,000 live cells were collected and resuspended in cold resuspension buffer (RSB: 10 mM Tris-HCl, pH 7.4; 10 mM NaCl; 3 mM MgCl₂) supplemented with 0.1% NP-40, 0.1% Tween-20, and 0.01% digitonin to permeabilize the plasma membrane. Following lysis, nuclei were washed in RSB containing 0.1% NP-40 and subjected to tagmentation using the transposition mixture: 25 μL of 2× TD buffer, 2.5 μL of Tn5 transposase (100 nM final concentration), 16.5 μL of 1× PBS, 0.5 μL of 1% digitonin, 0.5 μL of 10% Tween-20, and 5 μL of nuclease-free water. After transposition, DNA was purified using the Zymo DNA Clean & Concentrator-5 Kit (Zymo Research, #D4014). Library amplification was carried out as per protocol (Corces et al., 2017). Amplified libraries were purified using AMPure XP beads (Beckman Coulter, #A63880) with a double-sided size selection to remove both primer dimers and large fragments (>1000 bp). Final libraries were quantified using the Qubit dsDNA High Sensitivity Assay Kit (Invitrogen, #Q32854), pooled at equimolar concentrations, and sequenced on a NovaSeq X platform with a 10B flow cell, generating 150 bp paired-end reads.

For data processing, read quality was assessed using FastQC, and summary reports were generated with MultiQC (v1.25.2). Paired-end reads were trimmed using Trim Galore, and only high-quality reads passing these steps were retained for downstream analysis. Reads were aligned to the *Mus musculus* reference genome (mm10) using Bowtie2 (v2.4.1). Unaligned reads, mitochondrial reads, and PCR duplicates were removed at this stage. Peak calling was performed with MACS3 (v3.0.3), using --nomodel and --nolambda parameters. Peaks overlapping with mouse ENCODE blacklist regions were excluded. Library quality was further assessed by calculating the Fraction of Reads in Peaks (FRiP), enrichment at promoter regions, fragment length distribution, and genome-wide peak distribution. Differential accessibility analysis was performed in R using the DiffBind package (v3.16.0). Regions were considered significantly differentially accessible with a false discovery rate (FDR) < 0.05 and an absolute log₂ fold change > 1. Genomic regions were annotated to nearby genes using GREAT (McLean et al., 2010), with the following settings: proximal = ±5 kb from transcription start site (TSS); distal = up to 1000 kb. Principal component analysis (PCA) and heatmaps were generated from normalized count data using DiffBind. Gene ontology (GO) enrichment analysis was performed using the PANTHER database. Transcription factor enrichment analysis was conducted using ChEA via the ShinyGO web tool (v0.82).

### Public ChIP-seq analysis

For Smad2 ChIP-seq, publicly available bigWig files from mESCs were obtained from the Gene Expression Omnibus (GEO) database (GSE240787, GSE186222, and GSM5983538) (Zhao et al., 2024). To integrate these datasets, bigWig files were aggregated using deepTools (v3.5.4) bigwigCompare with ‘--operation add’. To ensure equal weighting of each dataset, scale factors were calculated and applied using ‘--scaleFactors’, normalizing the signal based on the total signal of each individual bigWig file. Following aggregation, the merged bigWig track was converted to bedGraph format using bigWigToBedGraph (UCSC Genome Browser utilities, obtained on April 16, 2024). The resulting bedGraph file was then sorted and peak regions were subsequently identified from the sorted bedGraph file using MACS2 (v2.2.7.1) bdgpeakcall function with default parameters and ‘-c 7’.

For Pparg ChIP-seq, publicly available bigWig files (Pparg-HA and HA control) were obtained from the GEO database (GSE199426) and aggregated using deepTools (v3.5.4) bigwigCompare with ‘--operation add’. Pparg peaks were taken from Lefterova et al (GSM532740 on GEO database) (Lefterova et al., 2010) and lifted from mm8 to mm10 using the UCSC Genome Browser LiftOver web tool (http://genome.ucsc.edu/cgi-bin/hgLiftOver).

### Public scRNA-seq analysis

Single-cell RNA sequencing (scRNA-seq) data was obtained from the GEO database (GSE241462), using the developmental time points E4.5, EDG7.5, and EDG9.5. Analyses were performed using R (v.4.4.1) with the Seurat (v5.2.0) package.

Briefly, raw count data for each time point were loaded and reformatted into gene by cell matrices. Initial quality control was performed by filtering cells with fewer than 500 or more than 10,000 detected genes, and those with over 20% mitochondrial gene content. Processed individual datasets were then normalized, variable features identified, and data scaled. Principal Component Analysis (PCA) was performed for dimensionality reduction. Subsequently, cells were clustered using graph-based clustering (based on the first 20 principal components) and Uniform Manifold Approximation and Projection (UMAP) was used for two-dimensional visualization (also utilizing the first 20 principal components). Individual datasets were subsequently merged into a single Seurat object for integrated analysis. The combined object underwent re-normalization, re-identification of variable features, scaling, and PCA (utilizing the top 10 principal components). Clustering was then performed using graph-based methods with a resolution of 0.5 (based on the first 10 principal components), and Uniform Manifold Approximation and Projection (UMAP) was used for two-dimensional visualization (also using the first 10 principal components).

Epiblast cell clusters were identified within the integrated dataset based on the expression of known markers (*Sox2* and *Nanog* genes). For subsequent analyses focusing on the epiblast lineage, data were restricted to cells within these identified clusters. All gene signatures were obtained from the Molecular Signatures Database (MSigDB, https://www.gsea-msigdb.org/gsea/msigdb/) and scores were compiled by calculating the mean z-score-normalized expression levels of the relevant signature genes for each cell.

### Data integration

For comparison between RNA-seq datasets, integration was performed by merging processed data at the gene level. Specifically, for the integration of publicly available embryo RNA-seq data (Fig. S3D), data obtained from GEO (GSE143494, GSE220804, GSE220804) and a published study (Boroviak et al., 2015) were batch-corrected by z-score normalization before merging. GSEA for public data was performed using the fGSEA package (v.1.26.0), as detailed in ‘RNA-seq library preparation and data analysis’.

For in-house *Smad2* Knockout (KO1 and KO2) RNA-seq data, differential expression analysis was first conducted individually for each KO line against the WT control. Given the high correlation observed between the two *Smad2* KO lines (Fig. S4C), KO1 and KO2 samples were grouped for a single differential analysis (comparing the combined KOs against WT), which was used throughout the study, unless otherwise stated.A similar merging principle was applied to in-house ATAC-seq data, where differential accessibility regions (DARs) correspond to the combined *Smad2* KO1 and KO2 versus WT comparison, unless explicitly stated otherwise.

When integrating RNA-seq and ATAC-seq data, comparisons were performed at the gene level after assigning ATAC peaks to corresponding genes using Genomic Regions Enrichment of Annotations Tool (GREAT) as detailed in the ‘ATAC-Seq library preparation and data analysis’ section. For ATAC-seq and ChIP-seq integration, two main approaches were used: (1) heatmaps of Pparg (or control HA) signal were generated over DAR regions using deepTools to visualize enrichment patterns; or (2) for assessing the overlap between DARs and Pparg target regions, respective peaks were first assigned to genes, and then the comparison was performed at the gene level using Venn diagrams.

### Data availability

Sequencing data have been deposited into the NCBI Gene Expression Omnibus (GEO) repository under accession number GSE303717.

### Protein extraction and western blotting

Cells were lysed in RIPA buffer (150 mM NaCl, 1% Nonidet P-40 Substitute, 0.5% sodium deoxycholate, 0.1% SDS, and 50 mM Tris-HCl, pH 8.0) supplemented with protease and phosphatase inhibitors (Life Technologies, #78442). Lysates were incubated on ice for 30 minutes, vortexed briefly, and centrifuged for 10 minutes at 20,000 × g. Supernatants were collected, and proteins were denatured at 95 °C for 10 minutes in Laemmli sample buffer. Proteins were separated on 4–15% Mini-Protean TGX SDS–PAGE gels (Bio-Rad, #4561083) and transferred to nitrocellulose membranes (Bio-Rad, #1704271). Membranes were blocked in 5% milk prepared in Tris-buffered saline with 0.1% Tween-20 (TBS-T) for 1 hour at room temperature, then incubated overnight at 4 °C with primary antibodies. The following day, membranes were washed three times for 10 minutes in TBS-T, incubated with HRP-conjugated secondary antibodies for 1 hour at room temperature, and washed again three times for 10 minutes. Signal detection was performed using ECL (Life Technologies, #32106) or Clarity Max (Bio-Rad, #1705062) chemiluminescent reagents. Images were acquired with Bio-Rad Image Lab Software. Band intensities were quantified using ImageJ and normalized to H3 levels. A full list of antibodies used and their dilution is provided in Supplementary Table 1.

### Immunofluorescence

Cells were plated on circular coverslips (EMS, #72222-01) and cultured as described above. At the time of collection, cells were fixed with 4% paraformaldehyde (Fisher Scientific, #AAJ61899-AP) for 10 minutes at room temperature, washed three times in PBS (10 minutes each), permeabilized with 0.1% Triton X-100 for 30 minutes at room temperature, and blocked with 1% bovine serum albumin (BSA) for 1 hour. Cells were then incubated overnight at 4 °C with primary antibodies diluted in BSA. The following day, cells were washed three times in PBS and incubated with fluorophore-conjugated secondary antibodies for 1 hour at room temperature. Nuclei were stained FxCycle^TM^ Violet Stain (Life Technologies, #F10347). After three final PBS washes, coverslips were mounted using a Epredia™ Immu-mount (Cat: 9990402) and imaged on a Nikon N2 Confocal microscope

BODIPY live-cell imaging was performed according to the manufacturer’s instructions (Life Technologies, #D3922). Briefly, cells were incubated with 10 μM BODIPY diluted in DMEM GlutaMAX for 30 minutes, protected from light. After incubation, cells were washed for 30 seconds in HEPES-buffered saline (HBS). Cells were then stained with 5μM Draq5 (Biostatus, #DR05500) diluted in HBS for 10 minutes to label nuclei and imaged immediately. Images were acquired using the BC43 benchtop confocal microscope (Andor).

Embryos were fixed in 4% paraformaldehyde for 15 minutes at room temperature. Permeabilization was performed using 0.5% Triton X-100 in PBS supplemented with 5% fetal bovine serum (FBS) for 15 minutes. Embryos were then blocked in PBS containing 2.5% bovine serum albumin (BSA) and 5% donkey serum for 1 hour at room temperature, followed by overnight incubation at 4 °C with primary antibodies diluted in blocking solution. The following day, embryos were washed three times for 10 minutes each in blocking solution and incubated with fluorophore-conjugated secondary antibodies for 1 hour at room temperature. After three additional washes, nuclei were stained with FxCycle^TM^ Violet Stain (Life Technologies, #F10347) in fresh blocking solution. Embryos were then transferred to 0.1% Tween20 in PBS and imaged using a Leica DMI6000 spinning disk confocal microscope. A complete list of antibodies used, along with their dilutions, is provided in Supplementary Table 1.

### Statistics and Reproducibility

Statistical analyses were performed using GraphPad Prism (v.9.3.1) or R (v.4.4.1). Data are presented as mean ± standard deviation (SD), as indicated. Data distribution was assumed to be normal but was not formally tested. For comparisons where normality could be assumed, two-tailed Student’s *t*-tests and one-way or two-way ANOVA with Dunnett’s multiple comparisons test were applied. Time series data were modeled using linear regression on log₂-transformed *y*-values, and *P*-values were derived from the interaction term between time and the categorical variable of interest. Embryo genotyping data were analyzed using the χ² test against expected Mendelian ratios. Gene Set Enrichment Analysis (GSEA) was performed using the fGSEA package in R, with adjusted *P*-values reported as indicated. Correlation between monoclonal lines was assessed using Pearson correlation coefficient (r) on log₂ fold-change. P-values for overlaps were calculated by two-sided hypergeometric test. For in vitro experiments, all replicates represent independent experiments in which a subpopulation of parental cells was randomly assigned to control or treatment groups; no formal randomization method was used. For ex vivo experiments, WT embryos from 6 litters were pooled, then randomly assigned to a group before treatment. For in vivo experiments, data were obtained from at least three embryos per genotype across three independent litters (except for *Smad2*^−/-^ for which no embryo was recovered following hormonal diapause). Randomization was not required for in vivo experimental design, as embryos were processed collectively and genotyped only after collection. Data collection and analysis were not performed blind to experimental conditions.

## Acknowledgements

We thank A. Bulut-Karslioglu, J. Wrana, A. Jurisicova, F. Lavial and members of the Santos lab for critical reading of the manuscript; members of the TCAG core at SickKids (Toronto) for next-generation sequencing; staff at the LTRI Flow Cytometry Facilities, and the TCP Animal Resources for colony management and Smad2 mouse line re-derivation. GF was supported by the PDF Banting fellowship and the Banting and Best Diabetes Centre Fellowship. EC was supported by a fellowship “Chargé de recherches” from the Belgian Fonds de la Recherche Scientifique (FNRS, CR40011342). This work was supported by Canada 150 Research Chair in Developmental Epigenetics, Anne & Max Tanenbaum Chair in Molecular Medicine at Mount Sinai and CIHR Project Grant 178094 to M.R.-S.

## Author contributions

G.F., E.C. and M.R.-S. conceived the project. G.F. and E.C. designed and performed most of the experiments and interpreted data. B.M. and B.C. performed embryo imaging. S.A.M. assisted with ATAC-Seq analysis. E.J.R. expanded and provided Nodal, Smad2, Smad3 and Smad2/3 KO ESCs, shared the Smad2 KO mouse model, and provided critical feedback. G.F., E.C. and M.R.-S. wrote the manuscript with feedback from all authors. M.R.-S. supervised the project.

## Supplementary Figure Legends

**Figure S1.**
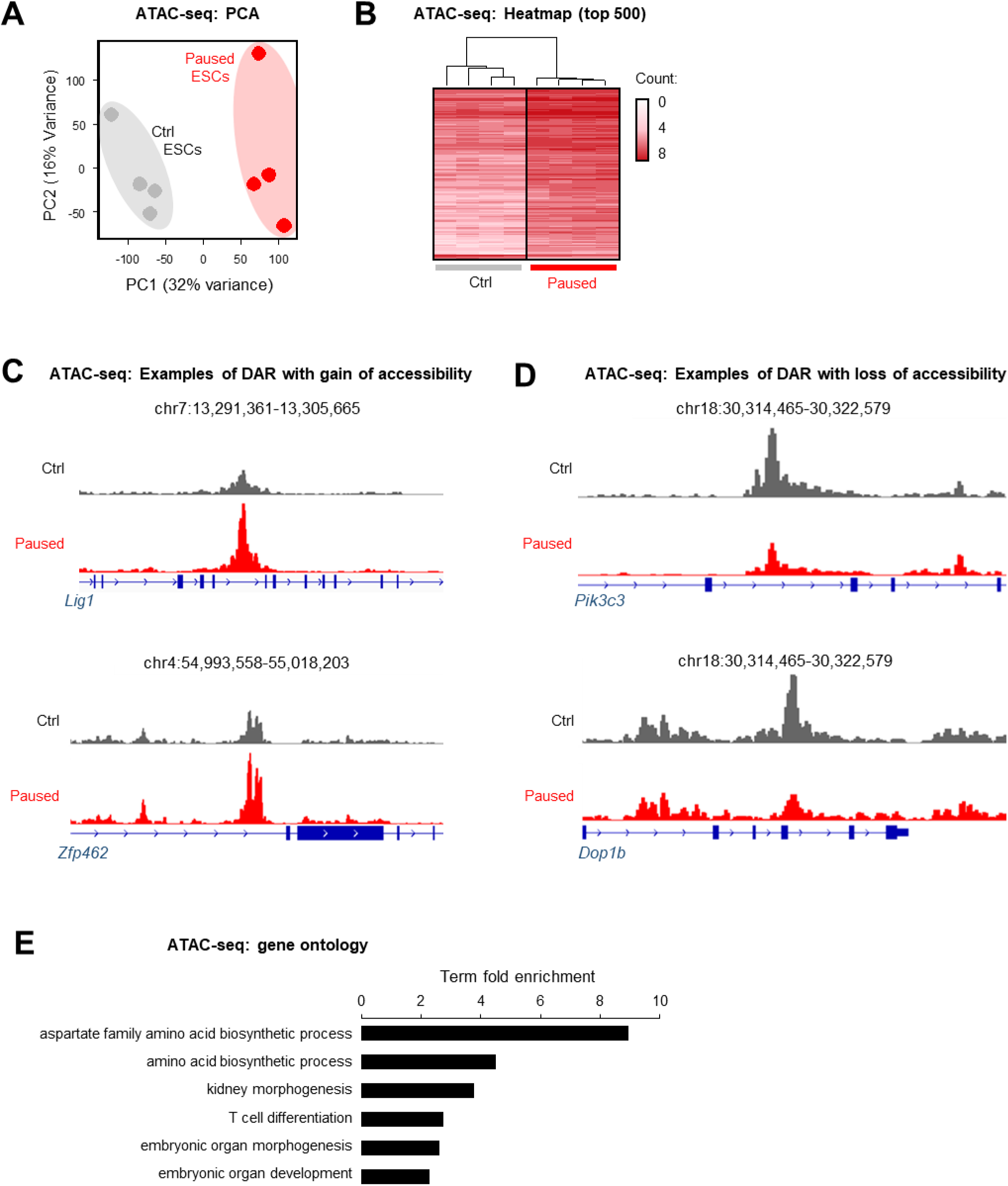
(related to Fig. 1) A. PCA plot of ATAC-seq data segregates paused and control ESC conditions. B. Heatmap of the top 500 differentially accessible sites between paused and control conditions, identified by ATAC-seq, with unsupervised hierarchical clustering. C-D. Examples of tracks showing increased (C) and decreased (D) genome accessibility (by ATAC-seq) in paused ESCs. E. Curated list of pathways enriched among sites with decreased accessibility (by ATAC-seq) using Panther Gene Ontology.

**Figure S2.**
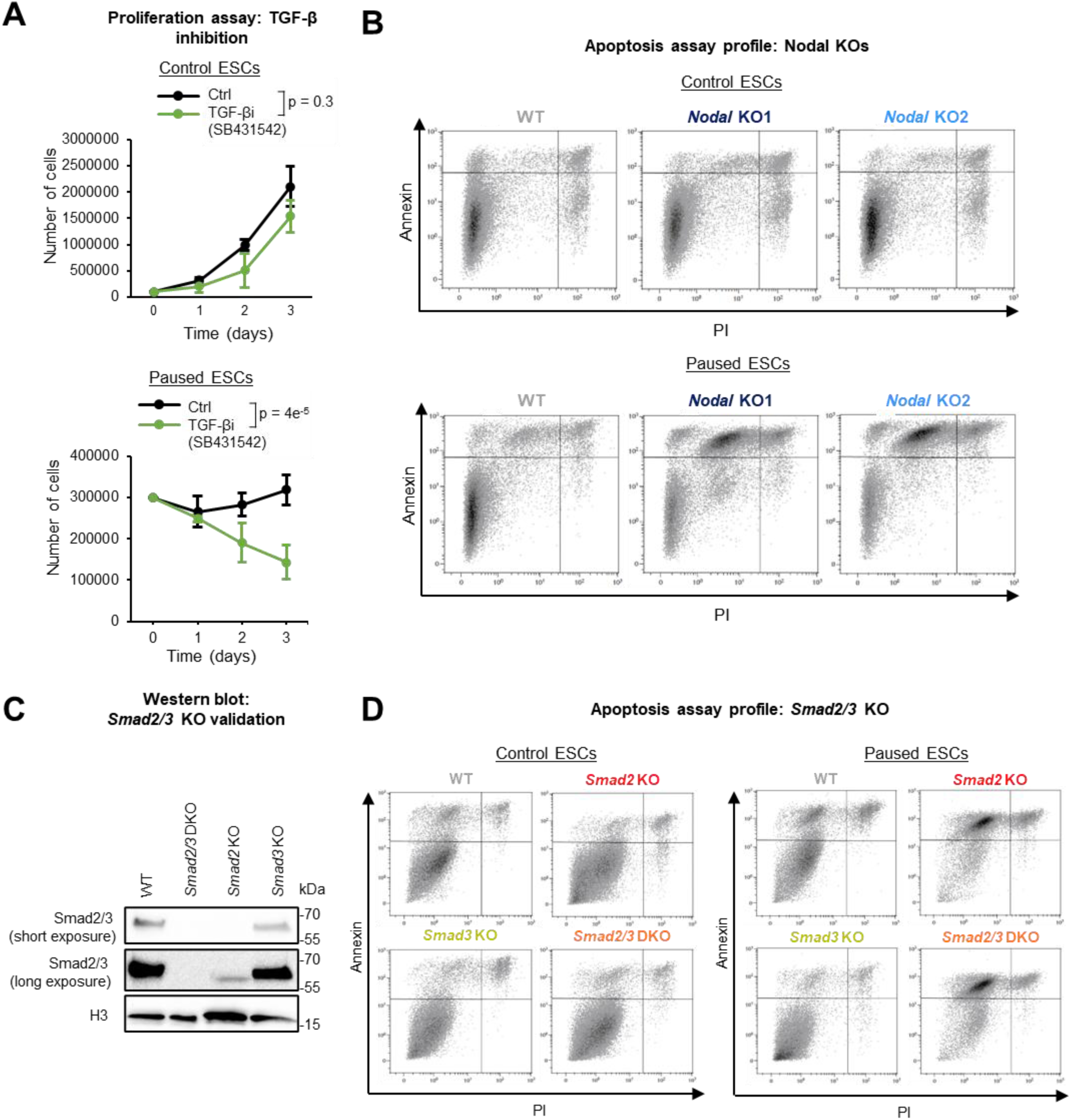
(related to Fig. 2) A. Proliferation assay of ESCs treated or not with 10µM TGF-β signaling inhibitor SB431542 in control (top) and paused (bottom) culture conditions. Data are presented as mean ± SD (n =3 biological replicates). B. Representative scatter plots of the apoptosis assays, as shown in Fig. 2B. C. Validation of *Smad2/3* DKO, *Smad2* KO, and *Smad3* KO in ESCs by western blot analysis, with histone H3 as loading control. D. Representative scatter plots of the apoptosis assays, as shown in Fig. 2F. P-values by linear regression test with interaction (A).

**Figure S3.**
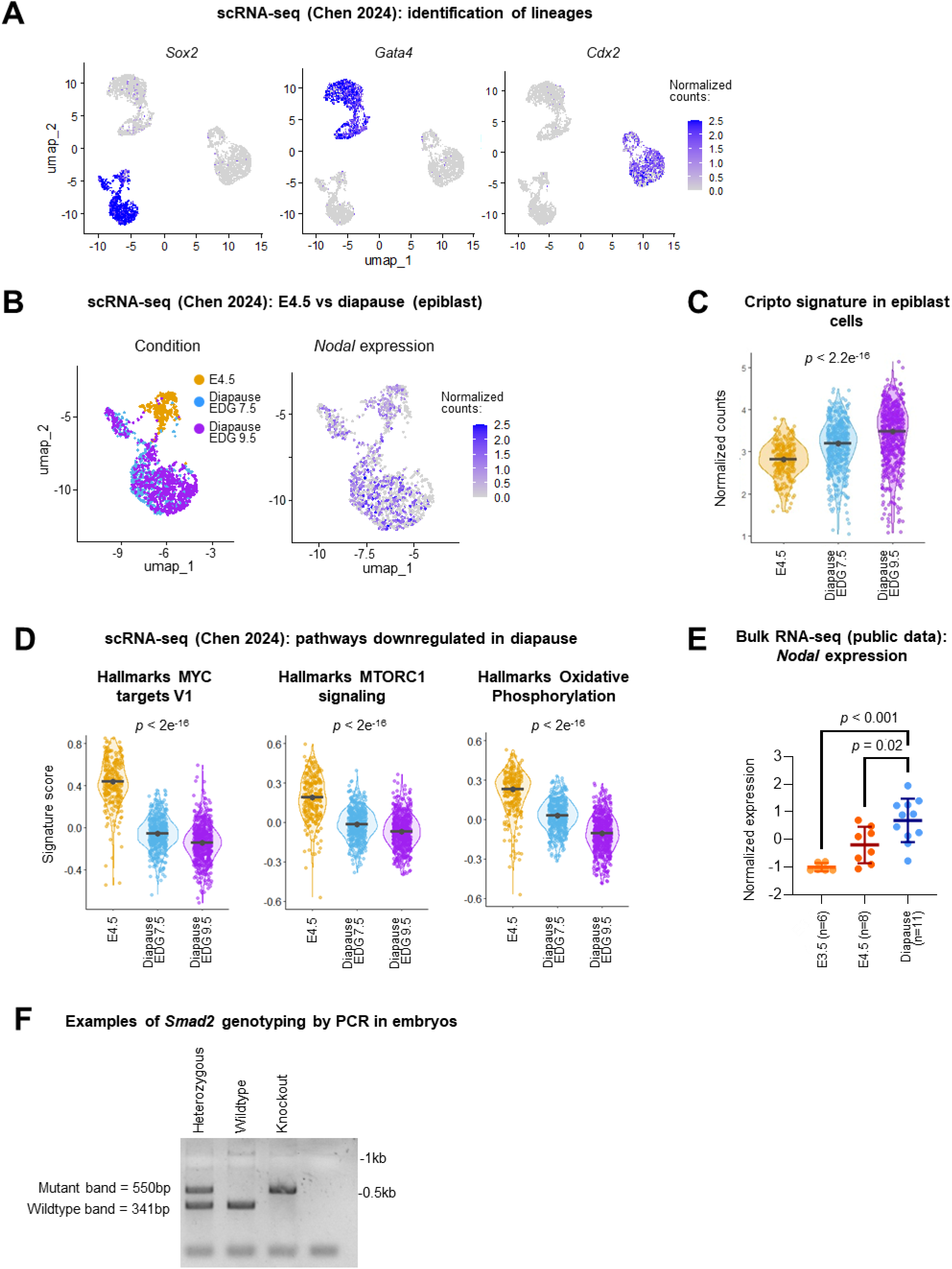
(related to Fig. 3) A. UMAP of diapaused (EDG7.5 and EDG9.5) and control blastocysts (E4.5) from public scRNA-seq data reveals three main clusters. Expression of *Sox2*, *Gata4*, and *Cdx2* identifies the epiblast, primitive endoderm, and trophectoderm lineages, respectively. B. Expression of *Nodal* among epiblast cells across all embryo conditions. C. Level of Cripto expression is increased in diapaused epiblast cells. D. Diapaused embryos exhibit the expected repression of MYC and mTORC1 signaling, and oxidative phosphorylation. E. Bulk RNA-seq data from four different studies (GSE220804) (Boroviak et al., 2015; Chen et al., 2024; Hussein et al., 2020) were batch normalized by z-score transformation and merged. Expression of *Nodal* is increased in diapaused embryos compared to control embryos (E3.5 or E4.5). F. Example of PCR genotyping of embryos resulting from *Smad2*^+/−^ crossing, representative of all genotyping performed in this study [n(*Smad2*^+/+^) = 13, n(*Smad2*^+/−^) = 26, n(*Smad2*^−/−^) = 4]. P-values by one-way ANOVA (C-D), or one-way ANOVA with Dunnett’s multiple comparison test (E).

**Figure S4.**
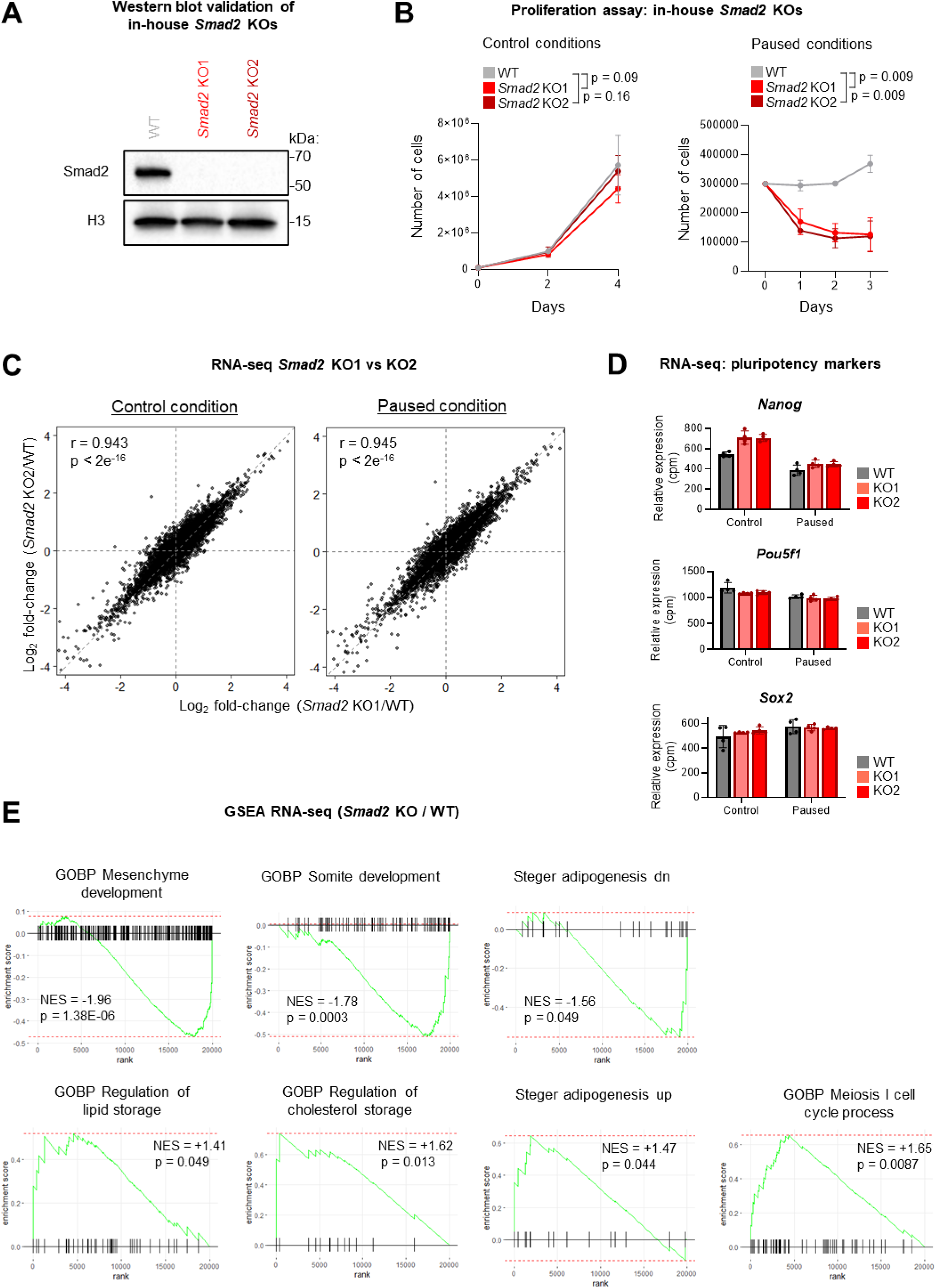
(related to Fig. 4) A. Validation of in-house *Smad2* KO1 and KO2 ESCs by western blot. Histone H3 as loading control. B. Functional validation of in-house *Smad2* KO1 and KO2 ESCs by proliferation assay. As seen in Fig. 2A, *Smad2* KO ESCs display a sharp decrease in cell number upon pausing, with moderate to no effect in control conditions. Data are presented as mean ± SD (n =2 biological replicates). C. *Smad2* KO1 and KO2 display highly consistent transcriptomic changes (compared to WT ESCs), both in control and paused conditions, as measured by RNA-seq. D. *Smad2* KO1 and KO2 show no differences in the expression of key pluripotent markers compared to WT, under either control or paused conditions. E. GSEA on RNA-seq data reveals lipid metabolism-associated pathways are enriched in paused *Smad2* KO ESCs, whereas various developmental pathways are decreased. P-values by linear regression test with interaction (B), Pearson correlation test (C), two-sided pre-ranked GSEA (E).

**Figure S5.**
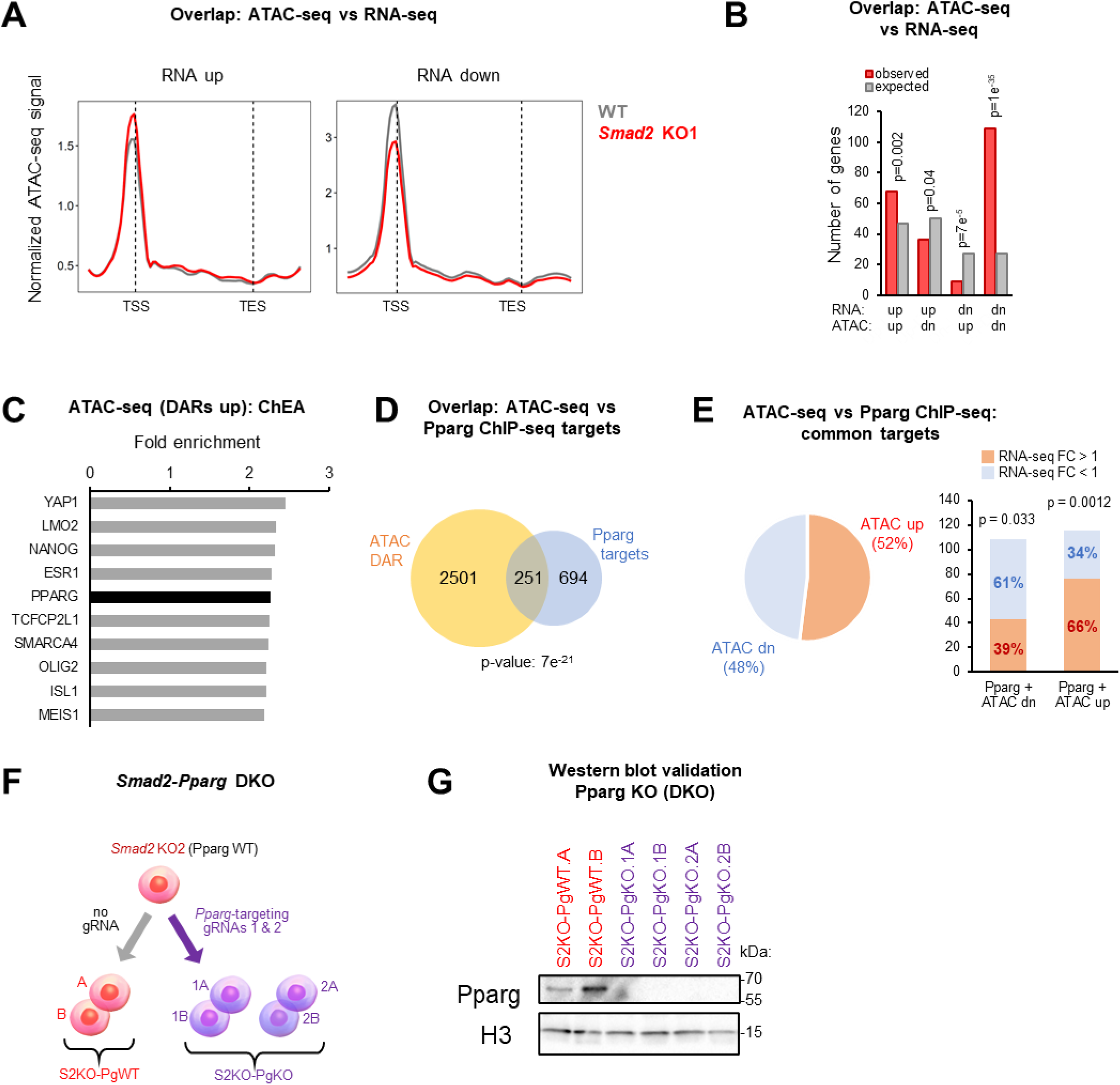
(related to Fig. 5) A. Metaprofiles of ATAC-seq signal from paused WT and *Smad2* KO1 ESCs, with genes upregulated in *Smad2* KO (‘RNA up’) showing increased promoter accessibility, while downregulated genes (‘RNA down’) exhibit a corresponding decrease. B. Bar plot showing the overlap between differentially expressed genes (by RNA-seq) and differentially accessible regions (by ATAC-seq) in *Smad2* KO paused ESCs. Genes with increased or decreased expression are significantly associated with corresponding changes in accessibility. C. ChIP-X Enrichment Analysis (ChEA) identifies enriched transcription factor motifs within DARs upregulated in paused *Smad2* KO ESCs. The top 10 enriched transcription factors, including PPARG, are shown. D. Venn diagram illustrates the significant overlap of genes associated with differentially accessible regions (DARs) in paused *Smad2* KO (by ATAC-seq) and Pparg ChIP-seq targets (Madsen et al., 2022). E. Comparison of ATAC-seq and RNA-seq changes on common ATAC-seq/Pparg targets (as defined in panel C). Left, pie chart showing the distribution of genes with increased (‘up’) and decreased (‘dn’) genome accessibility. Right, bar plot showing a significant association between changes in genome accessibility and gene expression (separated by RNA-seq fold-change >1 or <1). F. Schematic of *Smad2*–*Pparg* double knockout (DKO) generation by CRISPR-Cas9. Two *Pparg*-targeting gRNAs or no gRNA (empty vector) were introduced into *Smad2* KO2 ESCs, referred to as ‘S2KO-PgKO’ (1 and 2) and ‘S2KO-PgWT’, respectively. Two clones per condition (A and B) were selected for further analysis. G. Validation of in-house *Smad2-Pparg* DKO ESC clones. Pparg protein levels were assessed by western blot, with histone H3 as loading control. P-values by hypergeometric test for observed overlap (B), two-sided hypergeometric test (D), or two-sided one-proportion test (E).

## Bibliography

Aragón, E., Wang, Q., Zou, Y., Morgani, S.M., Ruiz, L., Kaczmarska, Z., Su, J., Torner, C., Tian, L., Hu, J., Shu, W., Agrawal, S., Gomes, T., Márquez, J.A., Hadjantonakis, A.-K., Macias, M.J., Massagué, J., 2019. Structural basis for distinct roles of SMAD2 and SMAD3 in FOXH1 pioneer-directed TGF-β signaling. Genes Dev 33, 1506–1524. 10.1101/gad.330837.119

Arena, R., Bisogno, S., Gąsior, Ł., Rudnicka, J., Bernhardt, L., Haaf, T., Zacchini, F., Bochenek, M., Fic, K., Bik, E., Barańska, M., Bodzoń-Kułakowska, A., Suder, P., Depciuch, J., Gurgul, A., Polański, Z., Ptak, G.E., 2021. Lipid droplets in mammalian eggs are utilized during embryonic diapause. Proc Natl Acad Sci U S A 118, e2018362118. 10.1073/pnas.2018362118

Baas, R., Sijm, A., van Teeffelen, H.A.A.M., van Es, R., Vos, H.R., Marc Timmers, H.T., 2016. Quantitative Proteomics of the SMAD (Suppressor of Mothers against Decapentaplegic) Transcription Factor Family Identifies Importin 5 as a Bone Morphogenic Protein Receptor SMAD-specific Importin. J Biol Chem 291, 24121–24132. 10.1074/jbc.M116.748582

Barcellos-Hoff, M.H., Gulley, J.L., 2023. Molecular Pathways and Mechanisms of TGFβ in Cancer Therapy. Clinical Cancer Research 29, 2025–2033. 10.1158/1078-0432.CCR-21-3750

Bertero, A., Brown, S., Madrigal, P., Osnato, A., Ortmann, D., Yiangou, L., Kadiwala, J., Hubner, N.C., de Los Mozos, I.R., Sadée, C., Lenaerts, A.-S., Nakanoh, S., Grandy, R., Farnell, E., Ule, J., Stunnenberg, H.G., Mendjan, S., Vallier, L., 2018. The SMAD2/3 interactome reveals that TGFβ controls m6A mRNA methylation in pluripotency. Nature 555, 256–259. 10.1038/nature25784

Boroviak, T., Loos, R., Lombard, P., Okahara, J., Behr, R., Sasaki, E., Nichols, J., Smith, A., Bertone, P., 2015. Lineage-Specific Profiling Delineates the Emergence and Progression of Naive Pluripotency in Mammalian Embryogenesis. Dev Cell 35, 366–382. 10.1016/j.devcel.2015.10.011

Brown, J.A., Yonekubo, Y., Hanson, N., Sastre-Perona, A., Basin, A., Rytlewski, J.A., Dolgalev, I., Meehan, S., Tsirigos, A., Beronja, S., Schober, M., 2017. TGF-β-Induced Quiescence Mediates Chemoresistance of Tumor-Propagating Cells in Squamous Cell Carcinoma. Cell Stem Cell 21, 650–664.e8. 10.1016/j.stem.2017.10.001

Bulut-Karslioglu, A., Biechele, S., Jin, H., Macrae, T.A., Hejna, M., Gertsenstein, M., Song, J.S., Ramalho-Santos, M., 2016. Inhibition of mTOR induces a paused pluripotent state. Nature 540, 119–123. 10.1038/nature20578

Chen, R., Fan, R., Chen, F., Govindasamy, N., Brinkmann, H., Stehling, M., Adams, R.H., Jeong, H.-W., Bedzhov, I., 2024. Analyzing embryo dormancy at single-cell resolution reveals dynamic transcriptional responses and activation of integrin-Yap/Taz prosurvival signaling. Cell Stem Cell 31, 1262–1279.e8. 10.1016/j.stem.2024.06.015

Chiu, W.T., Charney Le, R., Blitz, I.L., Fish, M.B., Li, Y., Biesinger, J., Xie, X., Cho, K.W.Y., 2014. Genome-wide view of TGFβ/Foxh1 regulation of the early mesendoderm program. Development 141, 4537–4547. 10.1242/dev.107227

Choy, L., Derynck, R., 2003. Transforming growth factor-beta inhibits adipocyte differentiation by Smad3 interacting with CCAAT/enhancer-binding protein (C/EBP) and repressing C/EBP transactivation function. J Biol Chem 278, 9609–9619. 10.1074/jbc.M212259200

Collignon, E., 2024. Unveiling the role of cellular dormancy in cancer progression and recurrence. Curr Opin Oncol 36, 74–81. 10.1097/CCO.0000000000001013

Collignon, E., Cho, B., Furlan, G., Fothergill-Robinson, J., Martin, S.-B., McClymont, S.A., Ross, R.L., Limbach, P.A., Ramalho-Santos, M., 2023. m6A RNA methylation orchestrates transcriptional dormancy during paused pluripotency. Nat Cell Biol 25, 1279–1289. 10.1038/s41556-023-01212-x

Corces, M.R., Trevino, A.E., Hamilton, E.G., Greenside, P.G., Sinnott-Armstrong, N.A., Vesuna, S., Satpathy, A.T., Rubin, A.J., Montine, K.S., Wu, B., Kathiria, A., Cho, S.W., Mumbach, M.R., Carter, A.C., Kasowski, M., Orloff, L.A., Risca, V.I., Kundaje, A., Khavari, P.A., Montine, T.J., Greenleaf, W.J., Chang, H.Y., 2017. An improved ATAC-seq protocol reduces background and enables interrogation of frozen tissues. Nat Methods 14, 959–962. 10.1038/nmeth.4396

Datto, M.B., Frederick, J.P., Pan, L., Borton, A.J., Zhuang, Y., Wang, X.-F., 1999. Targeted Disruption of Smad3 Reveals an Essential Role in Transforming Growth Factor β-Mediated Signal Transduction. Molecular and Cellular Biology 19, 2495–2504. 10.1128/MCB.19.4.2495

Deng, Z., Fan, T., Xiao, C., Tian, H., Zheng, Y., Li, C., He, J., 2024. TGF-β signaling in health, disease, and therapeutics. Signal Transduct Target Ther 9, 61. 10.1038/s41392-024-01764-w

Dhimolea, E., de Matos Simoes, R., Kansara, D., Al’Khafaji, A., Bouyssou, J., Weng, X., Sharma, S., Raja, J., Awate, P., Shirasaki, R., Tang, H., Glassner, B.J., Liu, Z., Gao, D., Bryan, J., Bender, S., Roth, J., Scheffer, M., Jeselsohn, R., Gray, N.S., Georgakoudi, I., Vazquez, F., Tsherniak, A., Chen, Y., Welm, A., Duy, C., Melnick, A., Bartholdy, B., Brown, M., Culhane, A.C., Mitsiades, C.S., 2021. An Embryonic Diapause-like Adaptation with Suppressed Myc Activity Enables Tumor Treatment Persistence. Cancer Cell 39, 240–256.e11. 10.1016/j.ccell.2020.12.002

Du, J., Wu, Y., Ai, Z., Shi, X., Chen, L., Guo, Z., 2014. Mechanism of SB431542 in inhibiting mouse embryonic stem cell differentiation. Cell Signal 26, 2107–2116. 10.1016/j.cellsig.2014.06.002

Finck, B.N., 2006. PGC-1 coactivators: inducible regulators of energy metabolism in health and disease. Journal of Clinical Investigation 116, 615–622. 10.1172/JCI27794

Fournier, T., Thérond, P., Handschuh, K., Tsatsaris, V., Evain-Brion, D., 2008. PPARgamma and early human placental development. Curr Med Chem 15, 3011–3024. 10.2174/092986708786848677

Fujiki, K., Kano, F., Shiota, K., Murata, M., 2009. Expression of the peroxisome proliferator activated receptor gamma gene is repressed by DNA methylation in visceral adipose tissue of mouse models of diabetes. BMC Biol 7, 38. 10.1186/1741-7007-7-38

Garcia-Ojalvo, J., Bulut-Karslioglu, A., 2023. On time: developmental timing within and across species. Development 150, dev201045. 10.1242/dev.201045

Gomis, R.R., Alarcón, C., Nadal, C., Van Poznak, C., Massagué, J., 2006. C/EBPbeta at the core of the TGFbeta cytostatic response and its evasion in metastatic breast cancer cells. Cancer Cell 10, 203–214. 10.1016/j.ccr.2006.07.019

Granier, C., Gurchenkov, V., Perea-Gomez, A., Camus, A., Ott, S., Papanayotou, C., Iranzo, J., Moreau, A., Reid, J., Koentges, G., Sabéran-Djoneidi, D., Collignon, J., 2011. Nodal cis-regulatory elements reveal epiblast and primitive endoderm heterogeneity in the peri-implantation mouse embryo. Developmental Biology 349, 350–362. 10.1016/j.ydbio.2010.10.036

Gross, B., Pawlak, M., Lefebvre, P., Staels, B., 2017. PPARs in obesity-induced T2DM, dyslipidaemia and NAFLD. Nat Rev Endocrinol 13, 36–49. 10.1038/nrendo.2016.135

Hernandez-Quiles, M., Broekema, M.F., Kalkhoven, E., 2021. PPARgamma in Metabolism, Immunity, and Cancer: Unified and Diverse Mechanisms of Action. Front Endocrinol (Lausanne) 12, 624112. 10.3389/fendo.2021.624112

Hiratsuka, D., Aikawa, S., Hirota, Y., Fukui, Y., Akaeda, S., Hiraoka, T., Matsuo, M., Osuga, Y., 2023. DNA Methylation and Histone Modification Are the Possible Regulators of Preimplantation Blastocyst Activation in Mice. Reprod Sci 30, 494–525. 10.1007/s43032-022-00988-x

Hussein, A.M., Wang, Y., Mathieu, J., Margaretha, L., Song, C., Jones, D.C., Cavanaugh, C., Miklas, J.W., Mahen, E., Showalter, M.R., Ruzzo, W.L., Fiehn, O., Ware, C.B., Blau, C.A., Ruohola-Baker, H., 2020. Metabolic Control over mTOR-Dependent Diapause-like State. Dev Cell 52, 236–250.e7. 10.1016/j.devcel.2019.12.018

Iyer, D.P., Moyon, L., Wittler, L., Cheng, C.-Y., Ringeling, F.R., Canzar, S., Marsico, A., Bulut-Karslioğlu, A., 2024. Combinatorial microRNA activity is essential for the transition of pluripotent cells from proliferation into dormancy. Genome Res 34, 572–589. 10.1101/gr.278662.123

Kumari, R., Irudayam, M.J., Al Abdallah, Q., Jones, T.L., Mims, T.S., Puchowicz, M.A., Pierre, J.F., Brown, C.W., 2021. SMAD2 and SMAD3 differentially regulate adiposity and the growth of subcutaneous white adipose tissue. FASEB J 35, e22018. 10.1096/fj.202101244R

Kutyavin, V.I., Chawla, A., 2019. BCL6 regulates brown adipocyte dormancy to maintain thermogenic reserve and fitness. Proc Natl Acad Sci U S A 116, 17071–17080. 10.1073/pnas.1907308116

Lakshmi, S.P., Reddy, A.T., Reddy, R.C., 2017. Transforming growth factor β suppresses peroxisome proliferator-activated receptor γ expression via both SMAD binding and novel TGF-β inhibitory elements. Biochemical Journal 474, 1531–1546. 10.1042/BCJ20160943

Lefterova, M.I., Steger, D.J., Zhuo, D., Qatanani, M., Mullican, S.E., Tuteja, G., Manduchi, E., Grant, G.R., Lazar, M.A., 2010. Cell-specific determinants of peroxisome proliferator-activated receptor gamma function in adipocytes and macrophages. Mol Cell Biol 30, 2078–2089. 10.1128/MCB.01651-09

Liu, W.M., Cheng, R.R., Niu, Z.R., Chen, A.C., Ma, M.Y., Li, T., Chiu, P.C., Pang, R.T., Lee, Y.L., Ou, J.P., Yao, Y.Q., Yeung, W.S.B., 2020. Let-7 derived from endometrial extracellular vesicles is an important inducer of embryonic diapause in mice. Sci Adv 6, eaaz7070. 10.1126/sciadv.aaz7070

Lowe, L.A., Yamada, S., Kuehn, M.R., 2001. Genetic dissection of nodal function in patterning the mouse embryo. Development 128, 1831–1843. 10.1242/dev.128.10.1831

MacLean Hunter, S., Evans, M., 1999. Non-surgical method for the induction of delayed implantation and recovery of viable blastocysts in rats and mice by the use of tamoxifen and Depo-Provera. Mol. Reprod. Dev. 52, 29–32. 10.1002/(SICI)1098-2795(199901)52:1<29::AID-MRD4>3.0.CO;2-2

Madsen, M.S., Broekema, M.F., Madsen, M.R., Koppen, A., Borgman, A., Gräwe, C., Thomsen, E.G.K., Westland, D., Kranendonk, M.E.G., Koerkamp, M.G., Hamers, N., Bonvin, A.M.J.J., Pittol, J.M.R., Natarajan, K.N., Kersten, S., Holstege, F.C.P., Monajemi, H., van Mil, S.W.C., Vermeulen, M., Kragelund, B.B., Cassiman, D., Mandrup, S., Kalkhoven, E., 2022. PPARγ lipodystrophy mutants reveal intermolecular interactions required for enhancer activation. Nat Commun 13, 7090. 10.1038/s41467-022-34766-9

Marsolier, J., Prompsy, P., Durand, A., Lyne, A.-M., Landragin, C., Trouchet, A., Bento, S.T., Eisele, A., Foulon, S., Baudre, L., Grosselin, K., Bohec, M., Baulande, S., Dahmani, A., Sourd, L., Letouzé, E., Salomon, A.-V., Marangoni, E., Perié, L., Vallot, C., 2022. H3K27me3 conditions chemotolerance in triple-negative breast cancer. Nat Genet 54, 459–468. 10.1038/s41588-022-01047-6

Massagué, J., Sheppard, D., 2023. TGF-β signaling in health and disease. Cell 186, 4007–4037. 10.1016/j.cell.2023.07.036

McGraw, M.S., Rajput, S.K., Daigneault, B.W., 2024. PPAR-gamma influences developmental competence and trophectoderm lineage specification in bovine embryos. Reproduction 167, e230334. 10.1530/REP-23-0334

Mesnard, D., Guzman-Ayala, M., Constam, D.B., 2006. Nodal specifies embryonic visceral endoderm and sustains pluripotent cells in the epiblast before overt axial patterning. Development 133, 2497–2505. 10.1242/dev.02413

Moghal, N., Li, Q., Stewart, E.L., Navab, R., Mikubo, M., D’Arcangelo, E., Martins-Filho, S.N., Raghavan, V., Pham, N.-A., Li, M., Shepherd, F.A., Liu, G., Tsao, M.-S., 2023. Single-Cell Analysis Reveals Transcriptomic Features of Drug-Tolerant Persisters and Stromal Adaptation in a Patient-Derived EGFR-Mutated Lung Adenocarcinoma Xenograft Model. J Thorac Oncol 18, 499–515. 10.1016/j.jtho.2022.12.003

Mullen, A.C., Wrana, J.L., 2017. TGF-β Family Signaling in Embryonic and Somatic Stem-Cell Renewal and Differentiation. Cold Spring Harb Perspect Biol 9, a022186. 10.1101/cshperspect.a022186

Nonninger, T.J., Mak, J., Gerisch, B., Ramponi, V., Kawamura, K., Ripa, R., Schilling, K., Latza, C., Kölschbach, J., Serrano, M., Antebi, A., 2025. A TFEB-TGFβ axis systemically regulates diapause, stem cell resilience and protects against a senescence-like state. Nat Aging 5, 1340–1357. 10.1038/s43587-025-00911-4

Park, C.B., DeMayo, F.J., Lydon, J.P., Dufort, D., 2012. NODAL in the Uterus Is Necessary for Proper Placental Development and Maintenance of Pregnancy1. Biology of Reproduction 86. 10.1095/biolreprod.111.098277

Park, C.B., Dufort, D., 2011. Nodal expression in the uterus of the mouse is regulated by the embryo and correlates with implantation. Biol Reprod 84, 1103–1110. 10.1095/biolreprod.110.087239

Park, S.-Y., Kim, M.-J., Park, S.-A., Kim, J.-S., Min, K.-N., Kim, D.-K., Lim, W., Nam, J.-S., Sheen, Y.Y., 2015. Combinatorial TGF-β attenuation with paclitaxel inhibits the epithelial-to-mesenchymal transition and breast cancer stem-like cells. Oncotarget 6, 37526–37543. 10.18632/oncotarget.6063

Ran, F.A., Hsu, P.D., Wright, J., Agarwala, V., Scott, D.A., Zhang, F., 2013. Genome engineering using the CRISPR-Cas9 system. Nat Protoc 8, 2281–2308. 10.1038/nprot.2013.143

Rehman, S.K., Haynes, J., Collignon, E., Brown, K.R., Wang, Y., Nixon, A.M.L., Bruce, J.P., Wintersinger, J.A., Singh Mer, A., Lo, E.B.L., Leung, C., Lima-Fernandes, E., Pedley, N.M., Soares, F., McGibbon, S., He, H.H., Pollet, A., Pugh, T.J., Haibe-Kains, B., Morris, Q., Ramalho-Santos, M., Goyal, S., Moffat, J., O’Brien, C.A., 2021. Colorectal Cancer Cells Enter a Diapause-like DTP State to Survive Chemotherapy. Cell 184, 226–242.e21. 10.1016/j.cell.2020.11.018

Renfree, M.B., Fenelon, J.C., 2017. The enigma of embryonic diapause. Development 144, 3199–3210. 10.1242/dev.148213

Ribeiro, E.S., Santos, J.E.P., Thatcher, W.W., 2016. Role of lipids on elongation of the preimplantation conceptus in ruminants. Reproduction 152, R115–126. 10.1530/REP-16-0104

Robertson, E.J., 2014. Dose-dependent Nodal/Smad signals pattern the early mouse embryo. Semin Cell Dev Biol 32, 73–79. 10.1016/j.semcdb.2014.03.028

Rüegg, A.B., Ulbrich, S.E., 2023. Review: Embryonic diapause in the European roe deer - slowed, but not stopped. Animal 17 Suppl 1, 100829. 10.1016/j.animal.2023.100829

Russo, M., Chen, M., Mariella, E., Peng, H., Rehman, S.K., Sancho, E., Sogari, A., Toh, T.S., Balaban, N.Q., Batlle, E., Bernards, R., Garnett, M.J., Hangauer, M., Leucci, E., Marine, J.-C., O’Brien, C.A., Oren, Y., Patton, E.E., Robert, C., Rosenberg, S.M., Shen, S., Bardelli, A., 2024. Cancer drug-tolerant persister cells: from biological questions to clinical opportunities. Nat Rev Cancer 24, 694–717. 10.1038/s41568-024-00737-z

Sabatino, L., Fucci, A., Pancione, M., Carafa, V., Nebbioso, A., Pistore, C., Babbio, F., Votino, C., Laudanna, C., Ceccarelli, M., Altucci, L., Bonapace, I.M., Colantuoni, V., 2012. UHRF1 coordinates peroxisome proliferator activated receptor gamma (PPARG) epigenetic silencing and mediates colorectal cancer progression. Oncogene 31, 5061–5072. 10.1038/onc.2012.3

Scognamiglio, R., Cabezas-Wallscheid, N., Thier, M.C., Altamura, S., Reyes, A., Prendergast, Á.M., Baumgärtner, D., Carnevalli, L.S., Atzberger, A., Haas, S., von Paleske, L., Boroviak, T., Wörsdörfer, P., Essers, M.A.G., Kloz, U., Eisenman, R.N., Edenhofer, F., Bertone, P., Huber, W., van der Hoeven, F., Smith, A., Trumpp, A., 2016. Myc Depletion Induces a Pluripotent Dormant State Mimicking Diapause. Cell 164, 668–680. 10.1016/j.cell.2015.12.033

Senft, A.D., Costello, I., King, H.W., Mould, A.W., Bikoff, E.K., Robertson, E.J., 2018. Combinatorial Smad2/3 Activities Downstream of Nodal Signaling Maintain Embryonic/Extra-Embryonic Cell Identities during Lineage Priming. Cell Reports 24, 1977–1985.e7. 10.1016/j.celrep.2018.07.077

Sirard, C., de la Pompa, J.L., Elia, A., Itie, A., Mirtsos, C., Cheung, A., Hahn, S., Wakeham, A., Schwartz, L., Kern, S.E., Rossant, J., Mak, T.W., 1998. The tumor suppressor gene Smad4/Dpc4 is required for gastrulation and later for anterior development of the mouse embryo. Genes Dev 12, 107–119. 10.1101/gad.12.1.107

Stötzel, M., Cheng, C.-Y., IIik, I.A., Kumar, A.S., Omgba, P.A., van der Weijden, V.A., Zhang, Y., Vingron, M., Meissner, A., Aktaş, T., Kretzmer, H., Bulut-Karslioğlu, A., 2024. TET activity safeguards pluripotency throughout embryonic dormancy. Nat Struct Mol Biol 31, 1625–1639. 10.1038/s41594-024-01313-7

Sun, H., Chen, Y., Yan, K., Shao, Y., Zhang, Q.C., Lin, Y., Xi, Q., 2023. Recruitment of TRIM33 to cell-context specific PML nuclear bodies regulates nodal signaling in MESCS. The EMBO Journal 42, e112058. 10.15252/embj.2022112058

Sun, Y., Zhang, L., Jiang, Z., 2024. The role of peroxisome proliferator-activated receptors in the regulation of bile acid metabolism. Basic Clin Pharmacol Toxicol 134, 315–324. 10.1111/bcpt.13971

Tan, C.K., Chong, H.C., Tan, E.H.P., Tan, N.S., 2012. Getting “Smad” about obesity and diabetes. Nutr Diabetes 2, e29. 10.1038/nutd.2012.1

Tremblay, K.D., Hoodless, P.A., Bikoff, E.K., Robertson, E.J., 2000. Formation of the definitive endoderm in mouse is a Smad2-dependent process. Development 127, 3079–3090. 10.1242/dev.127.14.3079

van der Weijden, V.A., Bulut-Karslioglu, A., 2021. Molecular Regulation of Paused Pluripotency in Early Mammalian Embryos and Stem Cells. Front Cell Dev Biol 9, 708318. 10.3389/fcell.2021.708318

van der Weijden, V.A., Stötzel, M., Iyer, D.P., Fauler, B., Gralinska, E., Shahraz, M., Meierhofer, D., Vingron, M., Rulands, S., Alexandrov, T., Mielke, T., Bulut-Karslioglu, A., 2024. FOXO1-mediated lipid metabolism maintains mammalian embryos in dormancy. Nat Cell Biol 26, 181–193. 10.1038/s41556-023-01325-3

Varlet, I., Collignon, J., Robertson, E.J., 1997. *nodal* expression in the primitive endoderm is required for specification of the anterior axis during mouse gastrulation. Development 124, 1033–1044. 10.1242/dev.124.5.1033

Waldrip, W.R., Bikoff, E.K., Hoodless, P.A., Wrana, J.L., Robertson, E.J., 1998. Smad2 Signaling in Extraembryonic Tissues Determines Anterior-Posterior Polarity of the Early Mouse Embryo. Cell 92, 797–808. 10.1016/S0092-8674(00)81407-5

Wang, X., Thiery, J.P., 2021. Harnessing Carcinoma Cell Plasticity Mediated by TGF-β Signaling. Cancers 13, 3397. 10.3390/cancers13143397

Weitlauf, H.M., Greenwald, G.S., 1968. Survival of blastocysts in the uteri of ovariectomized mice. J Reprod Fertil 17, 515–520. 10.1530/jrf.0.0170515

Yadav, H., Quijano, C., Kamaraju, A.K., Gavrilova, O., Malek, R., Chen, W., Zerfas, P., Zhigang, D., Wright, E.C., Stuelten, C., Sun, P., Lonning, S., Skarulis, M., Sumner, A.E., Finkel, T., Rane, S.G., 2011. Protection from obesity and diabetes by blockade of TGF-β/Smad3 signaling. Cell Metab 14, 67–79. 10.1016/j.cmet.2011.04.013

Zhao, B., Yu, X., Shi, J., Ma, S., Li, S., Shi, H., Xia, S., Ye, Y., Zhang, Y., Du, Y., Wang, Q., 2024. A stepwise mode of TGFβ-SMAD signaling and DNA methylation regulates naïve-to-primed pluripotency and differentiation. Nat Commun 15, 10123. 10.1038/s41467-024-54433-5

